# Invasive grass dominance over native forbs is linked to shifts in the bacterial rhizosphere microbiome

**DOI:** 10.1101/2021.01.07.425800

**Authors:** Marina L. LaForgia, Hannah Kang, Cassandra L. Ettinger

## Abstract

Rhizosphere microbiomes have received growing attention in recent years for their role in plant health, stress tolerance, soil nutrition, and invasion. Still, relatively little is known about how these microbial communities are altered under plant competition, and even less about whether these shifts are tied to competitive outcomes between native and invasive plants. We investigated the structure and diversity of rhizosphere bacterial and fungal microbiomes of native annual forbs and invasive annual grasses individually and in competition using high-throughput amplicon sequencing of the bacterial 16S rRNA gene and the fungal ITS region. We assessed how differentially abundant microbial families correlate to plant biomass under competition We find that bacterial diversity and structure differ between native forbs and invasive grasses, but fungal diversity and structure do not. Further, bacterial community structures under competition are distinct from individual bacterial community structures. We also identified five bacterial families that varied in normalized abundance between treatments and that were correlated with plant biomass under competition. We speculate that invasive grass dominance over these natives may be partially due to effects on the rhizosphere community, with changes in specific bacterial families potentially benefiting invaders at the expense of natives.

## Introduction

Plants and their associated soil microbial communities have coevolved complex dynamic relationships through time. The soil directly surrounding plant roots, known as the rhizosphere, is home to a diversity of microbes that play vital roles in plant health [1], nutrition [2], and stress tolerance [3, 4] and is distinct from the surrounding bulk soil [1,5–7]. Though rhizosphere composition is driven predominantly by abiotic factors [8–10], these communities can be highly host plant-specific [11, 12], with plants exhibiting strong local adaptations to their home microbial community [13–15]. Further, there is growing evidence that microbial communities vary in the presence of certain neighbors [16], and can even affect competitive outcomes between plants [17–19] with the potential to drive either coexistence or exclusion [20, 21].

Microbes can affect plant-host competition both directly via resource availability and indirectly through community interactions. For example, microbes can make nitrogen more readily available to plants [22, 23], which could preferentially benefit hosts with faster uptake rates [24, 25] thereby increasing their competitive success [26]. Plants may also indirectly compete for beneficial plant-growth-promoting microbes [7,27,28] or come into contact with novel pathogens through competition with a novel plant, such as an invader [29]. These interactions can lead to the loss of specialized microbes in the inferior competitor [30] and drive the resulting microbial community to resemble that of the dominant competitor [18, 19].

Shifts in the microbiome are a potentially important contributor to the dominance of invasive plants. Although there is strong evidence for local adaptation between plants and their home microbiome, invasive plants are relatively novel players, causing disruptions in important plant-microbe interactions by increasing soil microbial activity [31], reducing microbial biomass [32] and diversity [33], increasing nitrification rates [24], and changing microbiome composition [34, 35]. These plant-soil feedbacks can benefit the invader [35, 36], selectively harm natives [37], and/or benefit other invaders [38]. While it is known that invaders and natives can harbor distinct microbial communities, and that competition between host plants can affect microbiome composition (and vice versa), little is known of how changes in the abundance of specific microbial taxa may be tied to the competitive dominance of invaders and/or the competitive inferiority of natives.

Serpentine soils in California annual grasslands are characterized by high heavy metal content and low levels of essential plant nutrients, and as a result, are home to a unique community of native forbs. While some of these soils are rocky and shallow, other serpentine areas are deeper and more finely textured (“lush” serpentine) and therefore hold more water and nitrogen [39], making them more favorable to invasive annual grass establishment and growth. These fast-growing invasive grasses outcompete the less abundant and competitively inferior native forbs, contributing to their declines across California grasslands [40–42]. In native-dominated serpentine areas, microbes have been shown to increase seedling survival [43] and facilitate heavy metal tolerance [44]. With the invasion of grasses, microbiome changes may strengthen the advantage grasses have over natives if they recruit beneficial microbes away from natives or harbor novel microbes harmful to natives. Alternatively, locally adapted microbes may be harming invasive grasses, helping native forbs to persist in these areas.

We sought to understand how competition between native forbs and invasive grasses affects the bacterial and fungal rhizosphere microbiomes, and how microbes may be shaping competitive outcomes in this community. Our main questions were:

(Q1) Do microbiomes differ between invasive grasses and native forbs?

(Q2) Do microbiomes of grass-forb pairs (i.e. under competition) differ from the microbiomes of (a) invasive grasses and (b) native forbs?

(Q3) If so, are microbiomes of grass-forb pairs sourced more from invasive grasses, native forbs, or equally from both?

(Q4) Given compositional changes in the microbiome in grass-forb pairs, which specific microbial families are driving these changes?

(Q5) Are abundances of these microbial families correlated with plant performance in grass-forb pairs?

To answer these questions, we conducted a manipulative shade-house experiment in which forbs and grasses were grown in field-collected soil both individually and in grass-forb pairs. For each question, we looked at effects in both the bacterial and the fungal rhizosphere measured through high throughput amplicon sequencing, focusing our analysis on group-level results (i.e. forb, grass, and grass-forb pairs).

## Methods

### Experimental set-up

In spring and summer 2017, we collected soil and seed from a lush serpentine annual grassland at the University of California McLaughlin Natural Reserve [45] (N 38°52’, W 122°26’). Seeds were collected from >10 individuals each of six common native annual forb species (Fig. 1a; *Lasthenia californica, Clarkia purpurea, Agoseris heterophylla, Calycadenia pauciflora, Hemizonia congesta,* and *Plantago erecta)* and three dominant invasive annual grasses (Fig. 1b; *Avena fatua, Taeniatherum caput-medusae,* and *Bromus hordeaceus*). In December 2017, seeds were sown into physan-washed pots with a mixture of 65% field-collected soil and 35% sterile autoclaved sand to improve drainage. Each species was grown alone (one individual per pot, five replicate pots per species, 45 total pots: 30 forbs, 15 grasses) to assess the microbial communities of individual plant species. To assess the effects of competition between natives and invasives, each forb species was grown with each grass species in a pairwise factorial design for a total of 18 different species pairs (one individual of each grass and forb per pot, five replicate pots per combination, 90 total pots). We chose to focus on grass-forb pairs for our competition treatment as this treatment is the most relevant to understanding how microbial changes in the rhizosphere may contribute to invasive grass dominance. Due to plant mortality and a sample mix-up, final numbers were as follows: grass-forb pairs = 84, forbs = 27, grasses = 15. Pots were placed in a shade-house open to natural temperature variation in the UC Davis Orchard Park Greenhouse and soil moisture was maintained with an automatic drip irrigation system (Fig. 1c).

**Figure 1.**
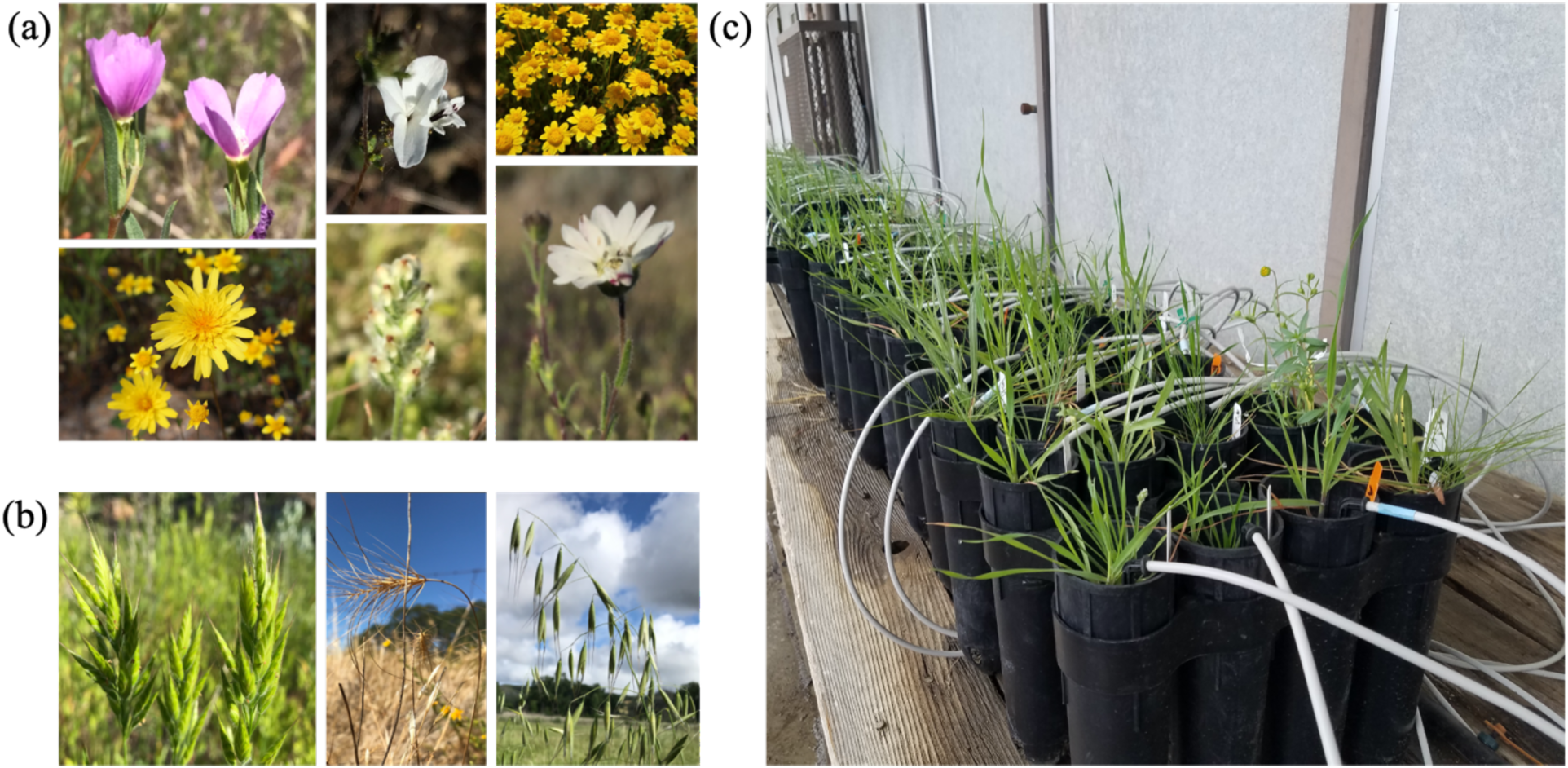
(a) Native annual forb species (from left to right, top to bottom): *Clarkia purpurea, Calycadenia pauciflora, Lasthenia californica, Agoseris heterophylla, Plantago erecta,* and *Hemizonia congesta.* (b) Invasive annual grasses (from left to right): *Bromus hordeaceus, Taeniatherum caput-medusae,* and *Avena fatua.* (c) Shade-house set-up. Photos of *C. purpurea*, *P. erecta, H. congesta*, *C. pauciflora*, *B. hordeaceus*, *T. caput-medusae*, and *A. fatua* by Paul Aigner. Remaining photos by authors.

### Sample collection

In April 2018, we sampled rhizosphere microbial communities by gently shaking off soil from plant roots, then submerging the roots in autoclaved nanopure water in a 50 mL conical tube and vortexing to obtain the rhizosphere soil similar to Edwards et al. [1]. Conical tubes were then centrifuged at 4000 g for 1 minute to obtain rhizosphere soil pellets. In grass-forb pairs, we sampled the joint rhizosphere microbiome, similar to Sun et al. [46], as roots were completely interwoven.

To assess how the microbial community may aid in invasive grass dominance over native forbs, we harvested, dried, and weighed aboveground plant biomass. Roots were not included due to the difficulty of identifying roots to species in the competition treatment. Previous work in this system has shown that competition with invasive grasses lowers fitness of native forbs through decreased seed production[41, 47]. While we were not able to measure seed production of plants in this study due to rhizosphere sampling prior to seed set, aboveground biomass in annual plants is often used as a surrogate for competitive ability [48, 49] and is well correlated with seed production [50, 51] and is therefore a reasonable estimate of plant performance.

### Molecular methods and sequence generation

Briefly, we extracted DNA from the 135 rhizosphere soil pellets using the MoBio PowerSoil DNA Extraction kit (MO BIO Laboratories, Inc., Carlsbad, CA, USA). Samples were then sent to the Integrated Microbiome Resource (IMR) at Dalhousie University to amplify and sequence the 16S rRNA gene using the 515FB-806RB primer set [52, 53] and ITS2 region using the ITS86F-ITS4R primer set [54]. The sequence reads generated for this 16S rRNA gene and ITS region amplicon project were deposited at Genbank under accession no. <u>PRJNA666893</u>.

### Sequence processing

Sequence data was processed using the DADA2 workflow [55] in R [56] to create Amplicon Sequence Variant tables (ASV). Chimeras, contaminants, mitochondria and chloroplasts were removed from these tables prior to further analysis. For questions 1-3, the 16S rRNA gene dataset was rarefied to 9434 reads per sample and the ITS region dataset was rarefied to 7557 reads per sample. These rarefaction levels were chosen, after examining rarefaction curves for saturation, based on the size of the sample with the smallest number of reads in order to retain the maximum number of samples for downstream analysis.

Further methodological details related to experimental set-up, sample collection and processing, sequence generation, and data processing prior to downstream analyses can be found in the Supplemental Materials.

### Data analysis (Q1-Q2)

To evaluate how rhizosphere communities varied in structure (beta diversity) between grasses, forbs, and grass-forb pairs, we conducted a Principal Coordinates Analysis (PCoA) on weighted UniFrac of each sample for rarefied 16S rRNA gene data and on Bray-Curtis dissimilarities for rarefied ITS region data. We tested for significant differences in mean centroids between groups (native forbs, invasive grasses, grass-forb pairs) using a permutational multivariate analysis of variance (PERMANOVA) and between mean dispersions using the betadisper function in vegan [57]. We then conducted post-hoc analyses with Benjamini-Hochberg p-value corrections [58]. We also investigated whether these groups varied compared to background soil (i.e. the community in the soil-sand mix at the start of the experiment) by performing PERMANOVA tests. Finally, we investigated the degree of species-level variation within both grasses and forbs by running PERMANOVA tests on each group separately to evaluate whether there was more variation within or between groups.

To assess how rhizosphere communities varied in alpha diversity between grasses, forbs and grass-forb pairs, we calculated Shannon diversity of each sample and conducted a Kruskal-Wallis test to test whether alpha diversity varied between these groups. We then conducted post-hoc analyses using a Dunn Test with Benjamini-Hochberg p-value corrections.

### Data analysis (Q3)

To evaluate whether a larger proportion of the joint microbiome community was predicted to be sourced from forbs or grasses, we used SourceTracker, a Bayesian source tracking classifier [59]. SourceTracker uses Gibbs sampling and Dirichlet distributions to estimate the proportional abundance of an ASV originating from provided source populations. First, we trained SourceTracker on the rarified ASV tables of forbs, grasses, and background soil and then tested the trained Bayesian model on the grass-forb pair joint microbiomes to estimate the proportion of ASVs originating from each of these sources. The model assumes that joint microbiomes contain a combination of colonists from known and unknown sources (i.e., any ASVs that are absent from the provided source datasets, and thus any sources not sampled here such as air, water, etc.) and estimates the fraction of ASVs detected in the joint microbiome that originated from grasses, forbs, background soil or unknown habitats. We used Wilcoxon signed-rank tests to assess differences in the SourceTracker predicted proportions between different sources for the bacterial and fungal communities, and then also tested for differences in the proportions of each individual source between the two communities.

### Data Analysis (Q4)

We used differential abundance analysis to identify microbial families whose abundance varied between groups in order to investigate which families were driving observed differences in community structure and composition. To do this, we first summed the raw read counts of each full data set (i.e. not rarefied), then used the DESeq2 R package [60] on counts to examine the log_2_fold change (i.e. differential abundance) of families between pairwise contrasts of groups (forbs, grasses, and grass-forb pairs). We chose to investigate differences at the family-level to maximize the potential for understanding the functional role of microbes while minimizing zero-inflation observed at higher resolutions. Only 1% of bacterial ASVs were unable to be taxonomically classified at the family level, while 49% of fungal ASVs could not be assigned to a family. Families with Bonferroni corrected *p*-values < 0.01 were classified as differentially abundant. In subsequent analyses, we focus on families that varied between both grass and forb individual microbiomes and between individual microbiomes and joint grass-forb pair microbiomes to highlight how baseline differences between groups are related to differences observed during competition.

### Data analysis (Q5)

To understand whether the abundances of these microbial families may be tied to the dominance of grasses over forbs, we first assessed how biomass varied between forbs and grasses both alone and in pairs, then we investigated the relationship between plant biomass in pairs and microbial family abundance. To compare biomass of forbs and grasses grown alone, we conducted a linear mixed effects model with biomass of plants grown alone as the response variable and functional group (grass or forb) as the predictor variable. Biomass was log-transformed to fit model assumptions and we included a plant species random intercept to account for species-level differences. To compare biomass of grasses and forbs in grass-forb pairs we ran a similar model on log biomass of plants in paired pots, again with a random intercept for each species. We explored other random effect structures that included a term accounting for non-independence of pots, however ultimately did not include this term as it did not improve the model based on the log-likelihood.

If grass dominance over forbs was controlled wholly by density dependent effects and competitive hierarchies rather than microbial interactions, we would expect an inverse relationship between grass biomass and forb biomass. To investigate the degree to which forb biomass was linked to grass biomass in pairs we used similar models with log-transformed forb biomass as the response variable and log-transformed grass biomass as the predictor variable and a random intercept for forb species.

To investigate the relationship between performance in grass-forb pairs and microbial abundance, we ran linear mixed effects models for each host group (grass and forb) and each differentially abundance microbial family with plant biomass from grass-forb pairs as the response variable and regularized log abundance of the focal family (calculated using the rld function in the DESeq2 package) as the predictor variable. Regularized log abundance (hereafter referred to as normalized abundance) is on the log2 scale and is normalized to library size to allow for direct comparison of microbial abundance across samples. We also included plant species as a random intercept.

Finally, we calculated the proportion of each differentially abundant family in the joint microbiome predicted to originate from either the individual forb or grass microbiomes by grouping ASVs into taxonomic families and then summarizing the fraction of ASVs within each family attributed to each possible source using SourceTracker.

## Results

### Bacterial microbiomes differ between invasive grasses and native forbs

Overall, the rhizosphere bacterial microbiome varied between invasive grasses and native forbs, but the fungal microbiome did not. The structure of forb bacterial microbiomes was significantly different from the structure of grass bacterial microbiomes (*p* < 0.001; Fig. 2). Further, bacterial Shannon diversity (Fig. 3) was significantly higher in forb microbiomes than in grass microbiomes (*p* = 0.035). Conversely, fungal communities showed marginally nonsignificant differences in structure (*p* = 0.07; Fig. S1) and no significant differences in Shannon diversity of fungal communities among treatments (*p* = 0.85; Fig. S2). Community structure was significantly different between background soil and all treatment groups for both the bacterial (*p <* 0.001, Fig. S3a) and fungal communities (*p* < 0.001, Fig. S3b). There were few species-level differences, with more variation between groups than between species within groups (Figs. S4, S5).

**Figure 2.**
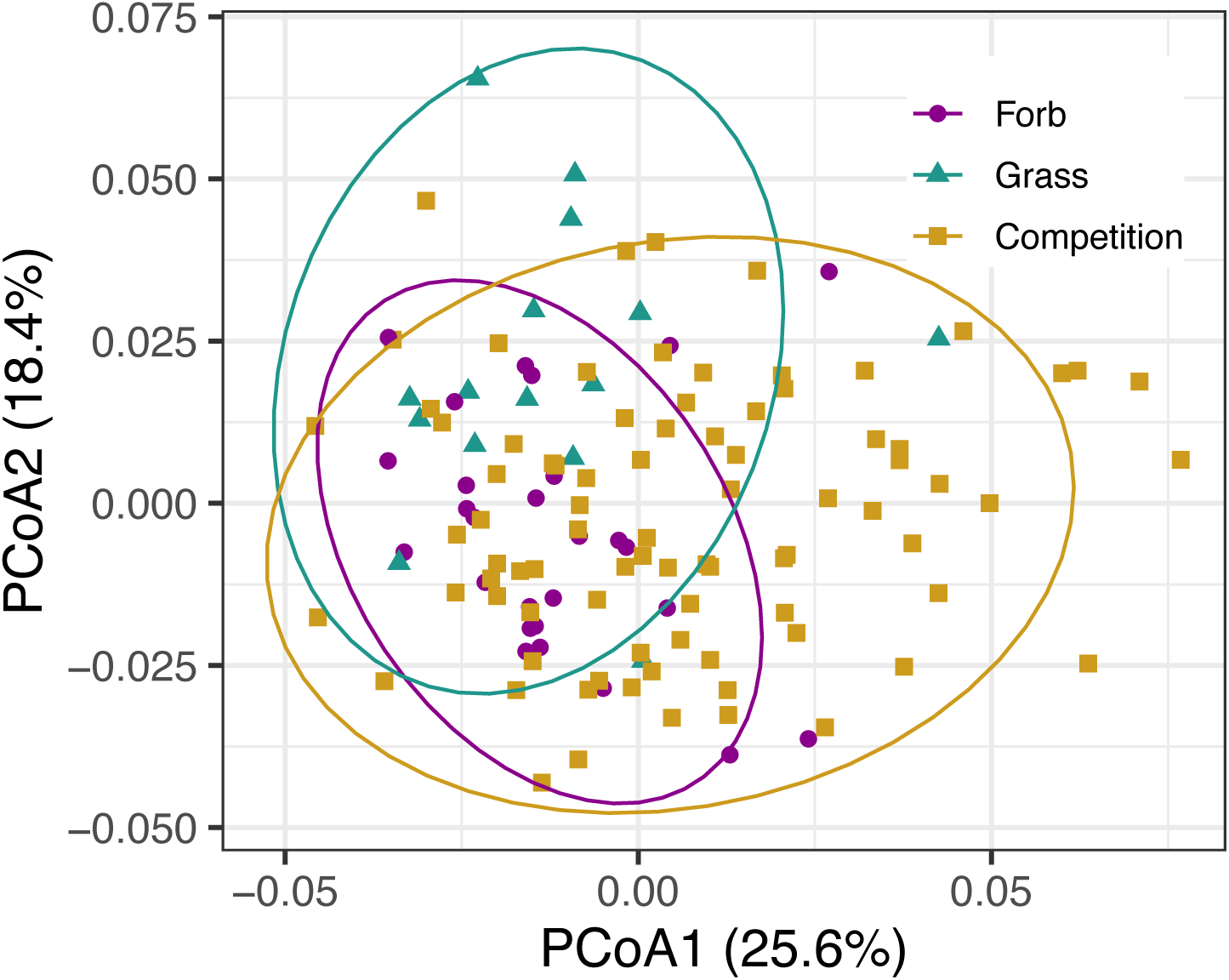
Bacterial microbiome structure differs between grasses and forbs, and shifts during competition. Principal coordinates analysis (PCoA) visualization of weighted UniFrac distances of bacterial communities associated with the rhizosphere. Points in the ordination are colored and represented by shapes based on the rhizosphere of forbs grown alone (purple circles), grasses grown alone (green triangles) and grass-forb pairs, i.e. competition (yellow squares). Ellipses represent the 95% confidence interval around the centroid of each group.

**Figure 3.**
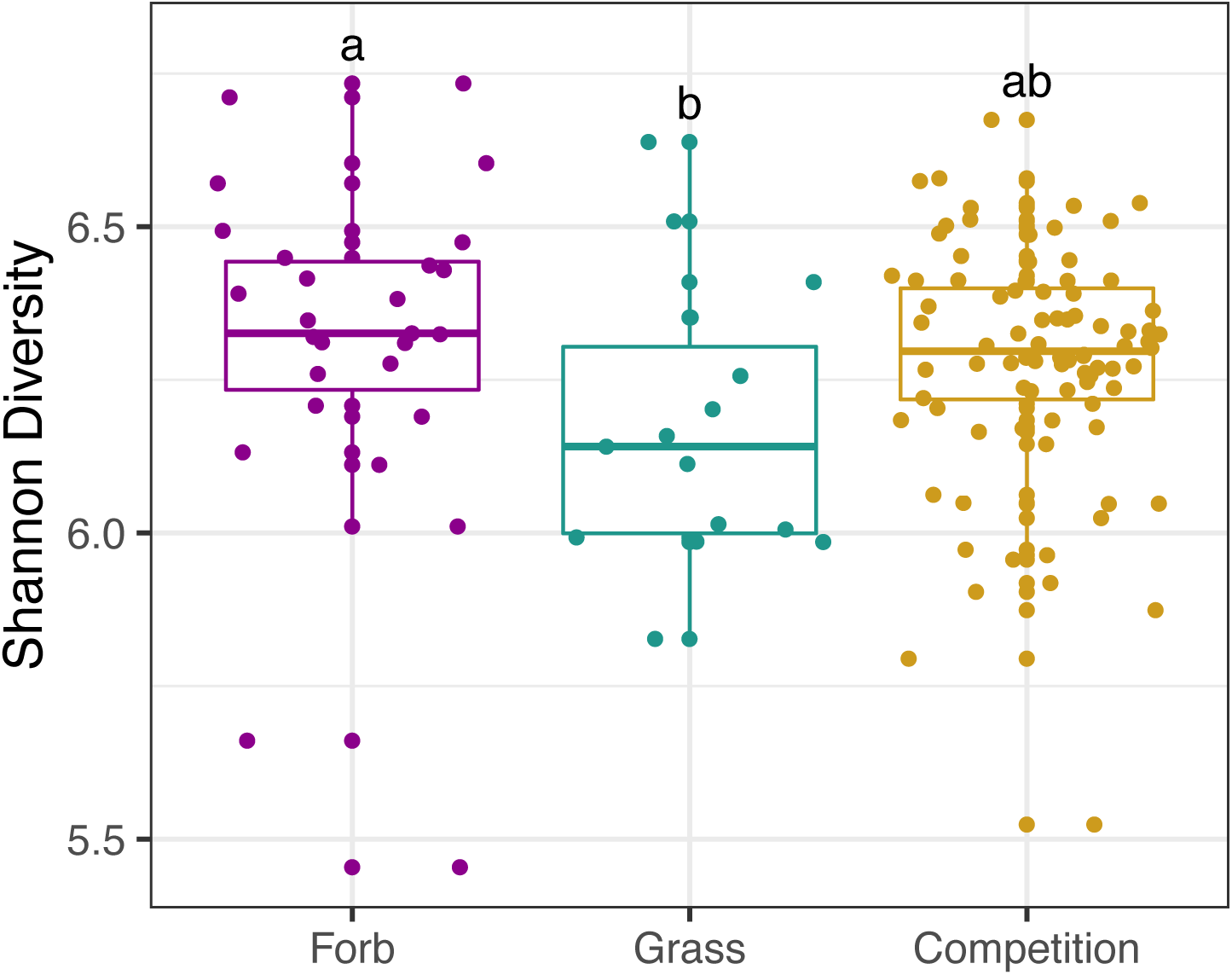
Shannon diversity of bacterial microbiomes is highest in forbs. Shannon diversity was used to assess alpha diversity for bacterial microbiomes for each treatment (forbs grown alone, grasses grown alone, competition). Comparisons that are significantly different from each other are notated by different letters (e.g. a vs. b).

### Bacterial microbiomes of grass-forb pairs differ from the microbiomes of grasses and forbs

Grass-forb pairs displayed significantly different joint bacterial microbiomes from microbiomes of both groups grown alone (forb: *p* < 0.001; grass: *p* < 0.001; Fig. 2). Grass-forb pair microbiomes also had marginally higher Shannon diversity than grass microbiomes (*p* = 0.059), but did not significantly differ from forb microbiomes (*p* = 0.292). As reported above, there were no significant differences found between treatments in the fungal microbiome, so we did not conduct post-hoc analyses to assess pairwise contrasts between treatment groups.

### Microbiomes of grass-forb pairs are sourced equally from invasive grasses and native forbs

For bacteria, SourceTracker identified similar fractions of ASVs in grass-forb pairs predicted to have originated from grasses and from forbs (*p* = 0.341, grass mean ± se: 40.34 ± 1.33%, forb: 42.13 ± 1.14%). Only relatively low proportions of the community came from background soil or unknown sources (background: 8.57 ± 0.48%, unknown: 8.96 ± 0.60%). For fungi, the fraction of the community predicted to originate from grasses was marginally higher than that from forbs (*p* = 0.066, grass: 38.88 ± 2.26%, forb: 32.65 ± 2.01%). Additionally, the fraction of the community originating from forbs was significantly lower than the fraction of the bacterial community originating from forbs (*p* < 0.001). A small proportion of the fungal community was predicted to derive from background soil (5.16 ± 0.67%). A much larger proportion of the fungal community was estimated to be from unknown sources (23.31 ± 2.11%), which was significantly higher than the contribution of unknown sources to the bacterial community (*p* < 0.001).

### Six bacterial and one fungal family displayed differential abundances between forbs, grasses, and pairs

Differential abundance analysis (DESeq2) revealed 9 bacterial families (out of a total of 302 bacterial families) (Fig. S6a) that differed in abundance between individual grass and forb microbiomes, with six of these families remaining differentially between joint grass-forb pair microbiomes and individual microbiomes: Methylophilaceae, Fibrobacteraceae, Clostridiaceae_1, Burkholderiaceae, Rhodocyclaceae, and Veillonellaceae (Fig. 4). Of these six bacterial families, only Burkholderiaceae had an untransformed mean relative abundance of greater than one percent (11.69 ± 0.39%; Table S1). There were an additional 10 bacterial families that varied between pairs and individuals, but were not found to be differentially abundant between grasses and forbs grown alone (Fig. S7).

**Figure 4.**
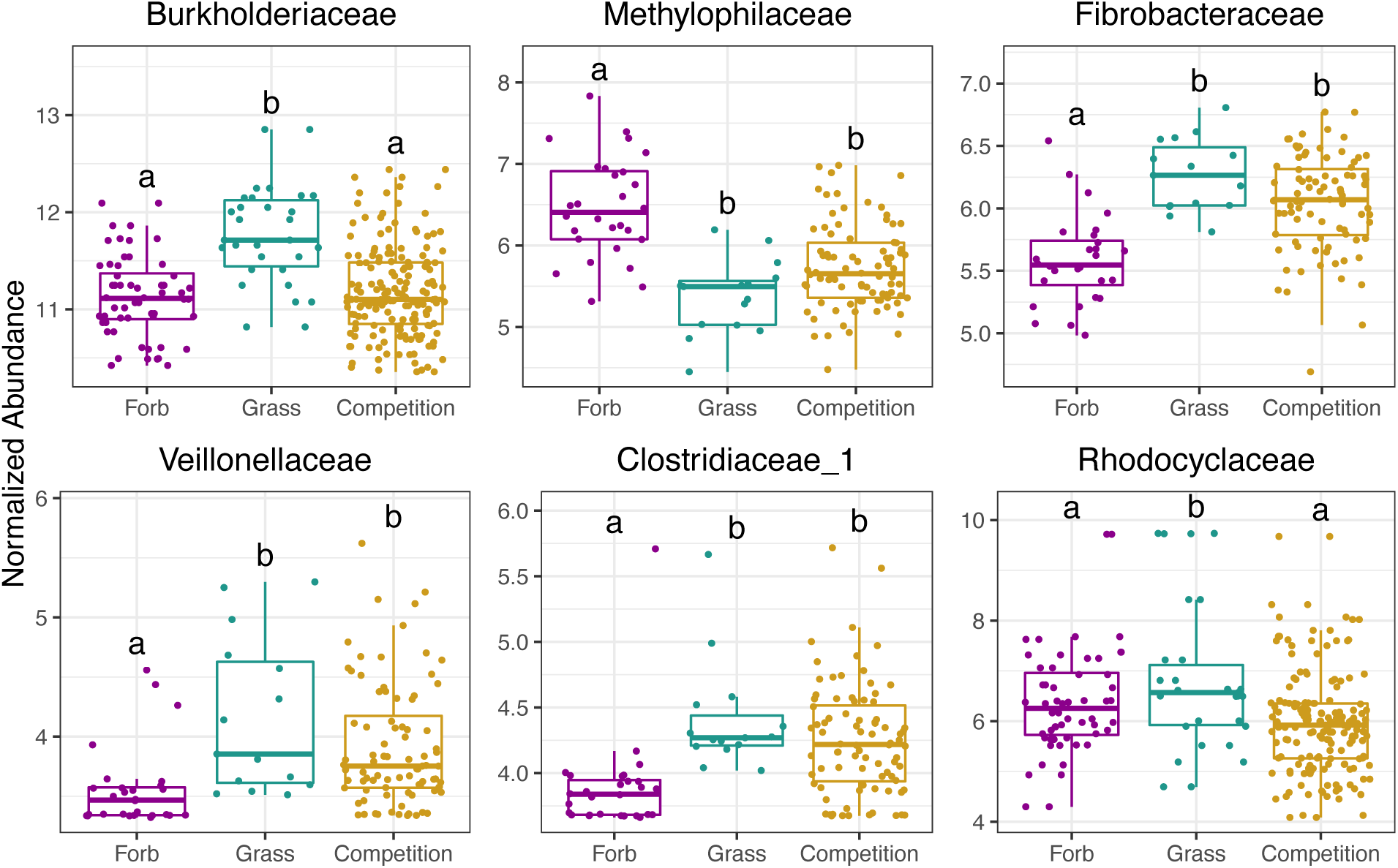
Bacterial families that are differentially abundant between grass and forb microbiomes, and between grass-forb pairs and either grass or forb microbiomes. Using DESeq2, bacterial families were identified whose abundance differed significantly between treatments (forbs grown alone, grasses grown alone, grown together in competition). Each plot shows the normalized abundance of families for each treatment and comparisons that are significant from each other are notated by different letters (e.g. a vs. b).

Methylophilaceae was the only family with higher normalized abundance in forbs than in grasses and their normalized abundance declined in pairs relative to forbs. Fibrobacteraceae, Clostridiaceae_1 and Veillonellaceae were all higher in normalized abundance in grasses than in forbs and increased in grass-forb pairs relative to forbs. Both Burkholderiaceae and Rhodocyclaceae were higher in grasses than in forbs, but decreased in grass-forb pairs relative to grasses.

Using DESeq2, we found only one fungal family (out of 231) that was differentially abundant between grasses and forbs, and between pairs and individuals (Tubeufiaceae) (Fig. S6b, Fig. S8). This family had higher normalized abundance in forbs than in grasses and subsequently lower normalized abundance in grass-forb pairs than in forbs. Two other families that did not vary between grasses and forbs showed variation between pairs and either grasses or forbs.

### Normalized abundances of key microbial families are correlated with grass dominance over forbs

When forbs and grasses were grown alone, grass and forb biomass did not significantly differ (*p* = 0.435), however in grass-forb pairs, grass biomass was significantly higher than forb biomass (*p* < 0.001), indicating that despite no inherent differences in biomass between groups at this stage, grasses outperformed forbs when grown together. We found no significant relationship between forb biomass and grass biomass in grass-forb pairs (est = -0.153, se = 0.102, *p* = 0.138), suggesting that decreases in forb biomass were not solely due to increases in grass biomass. Instead, individual plant performance in pairs was closely related to microbial abundance. Of the six families that differentially varied between grasses and forbs grown alone, and remained differentially abundant in pairs, all but Rhodocyclaceae displayed relationships with either forb or grass biomass in paired pots (Fig. 5, Table S2). Moreover, the four families that displayed negative correlations with forb biomass and/or positive correlations with grass biomass (Burkholderiaceae, Fibrobacteraceae, Clostridiaceae_1, and Veillonellaceae) had higher predicted proportions originating from grass microbiomes according to SourceTracker, while the one family that displayed a positive correlation with forb biomass and a negative correlation with grass biomass (Methylophilaceae) had a higher predicted proportion originating from forb microbiomes (Fig. 6).

**Figure 5.**
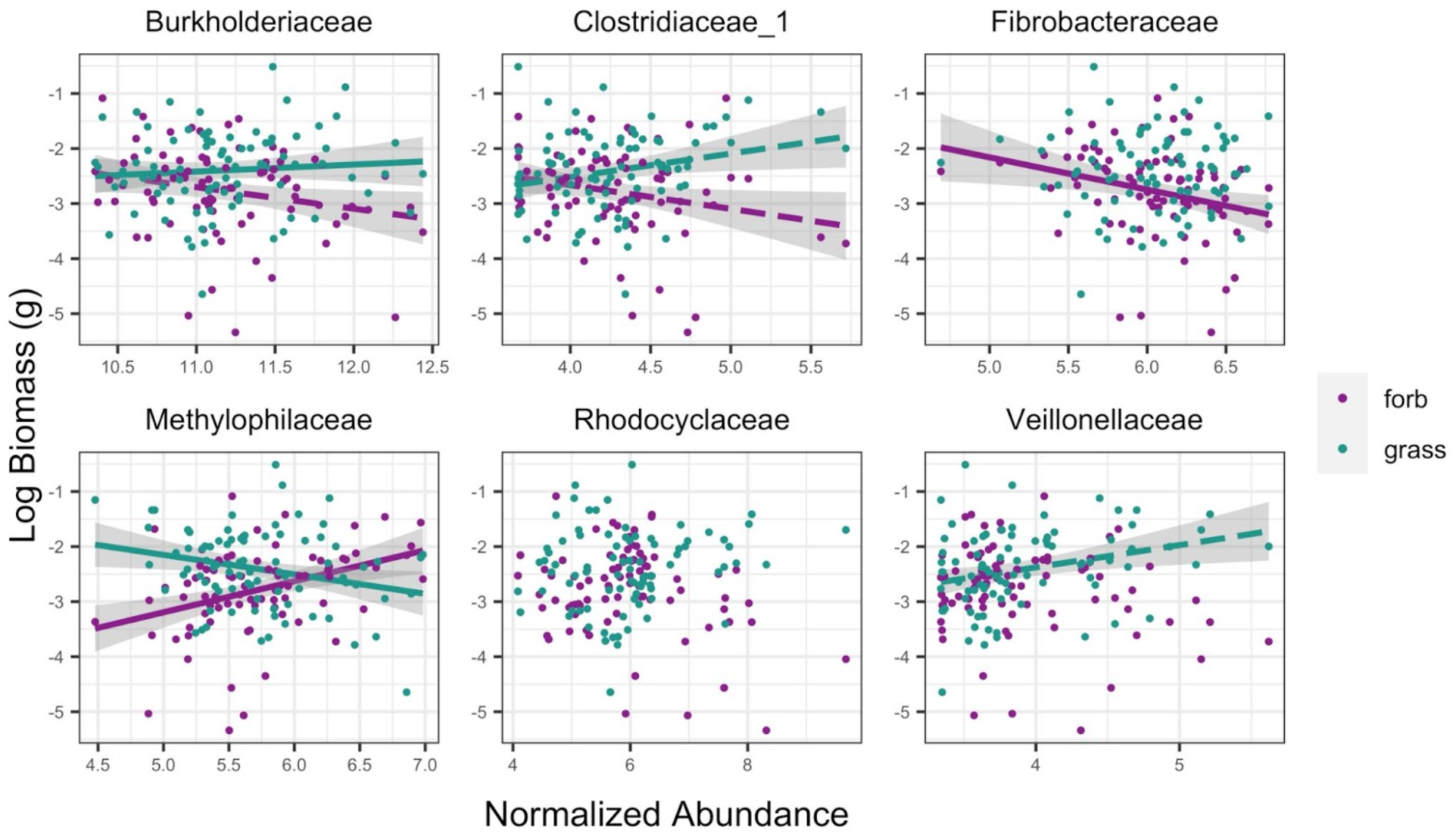
Bacterial family abundances correlated with forb (purple) and grass (green) biomass in pairs. Solid line indicates significance (*p* < 0.05), dashed line indicates marginal significance (*p* < 0.10). For regression results see Table S2.

**Figure 6.**
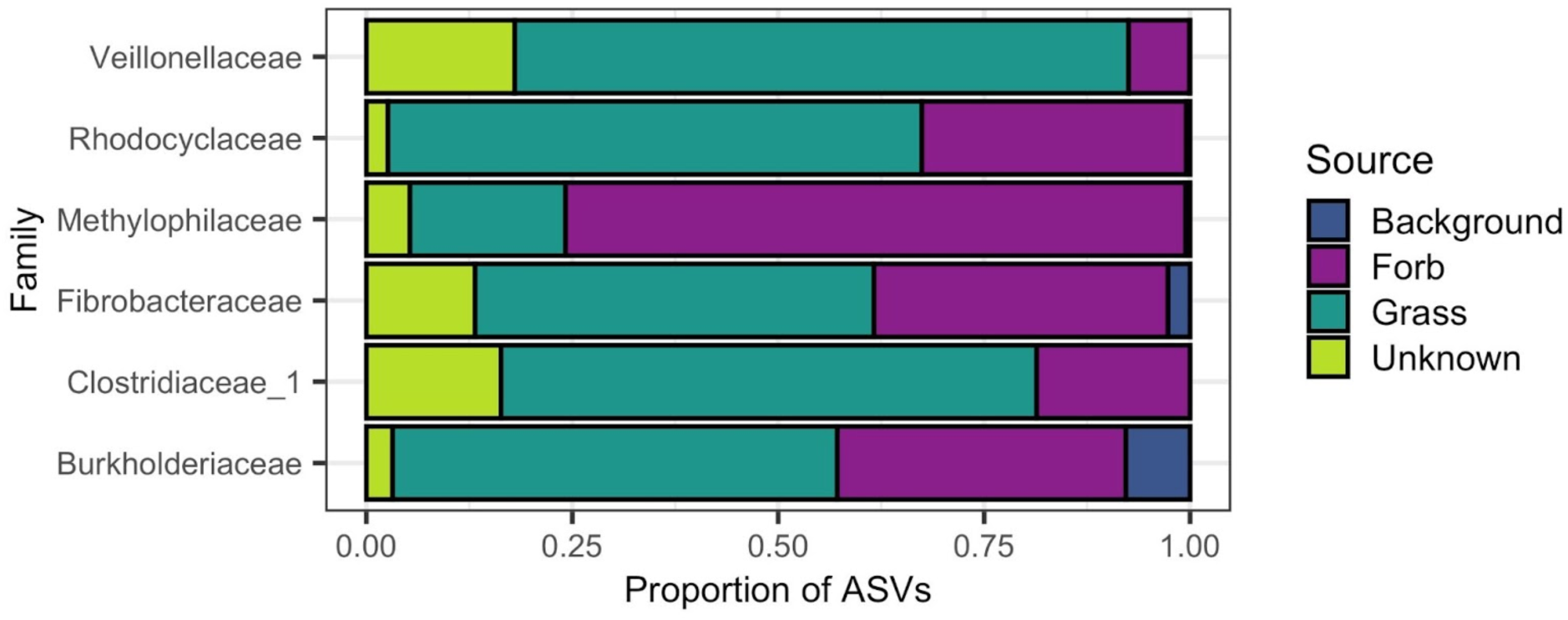
Proportion of ASVs predicted to colonize from each source for each of the six differentially abundant families. SourceTracker was used to predict whether ASVs in competition treatments originated from grasses (dark green), forbs (purple), the background soil mix used in the experiment (blue), or unknown habitats (light green) and then the results were summarized by grouping ASVs by taxonomic family. Families that were negatively correlated with forb performance and/or positively correlated with grass performance were predominantly sourced from grass. Methylophilaceae, the one family that showed a positive relationship with forb biomass under competition and a negative relationship with grass biomass under competition, was predominantly sourced from forbs.

The normalized abundance of Methylophilaceae was positively correlated with forb biomass in pairs, and negatively correlated with grass biomass in pairs, indicating that as the normalized abundance of this family decreased, grass performance increased and forb performance decreased. The normalized abundance of Fibrobacteraceae was negatively correlated with forb biomass in pairs, but was not correlated with grass biomass, indicating that as the normalized abundance of this family increased, forb performance decreased despite no change in grass performance. The normalized abundance of Clostridiaceae_1 was positively correlated with grass biomass, and marginally negatively correlated with grass biomass. The normalized abundance of Burkholderiaceae was positively correlated with grass biomass and marginally negatively correlated with forb biomass. Finally, the normalized abundance of Veillonellaceae was marginally positively correlated with grass biomass.

In the remaining 13 differentially abundant bacterial families, only Weeksellaceae, which did not vary between grasses and forbs, showed a significant relationship with biomass (Fig. S9, S10; Table S2). Similarly, only the fungal family Ceratobasidiaceae, which again did not vary between forbs and grasses, showed a significant relationship with biomass (Fig. S11, S12; Table S2).

## Discussion

The rhizosphere bacterial community of native forbs is structurally distinct from those of invasive grasses and has higher alpha diversity. Interactions between grasses and forbs here correlated with shifts in the bacterial rhizosphere towards communities dissimilar to both individual forb and grass microbiomes, with marginally higher diversity than grasses alone. Studies in similar annual grasslands have found little change in bacterial structure with increased invasive grass abundance [61–63] but invader dominance has been linked to decreased bacterial diversity in other systems [33, 64]. While we expected the microbiomes of grass-forb pairs to be a mixed community sourced from both forbs and grasses, we also expected the joint rhizosphere to be dominated by the microbes of the dominant competitor as in Hortal et al. [18] and Lozano et al. [19]. In partial support of this, we found that four out of the six families that varied both between forbs and grasses and between individuals and pairs came predominantly from grasses and increased in abundance relative to forbs, while the one family that was majority sourced from forbs saw a decrease in abundance. At the community level, however, we found that the novel assemblage of bacteria in pairs was sourced nearly equally from both groups. These results suggest that the invasions of grasses into native forb habitats may be associated with microbiome shifts in both groups; however, these changes may have more negative consequences for native forbs than invasive grasses.

Consistent with invasive annual grasses aiding in native forb declines across California annual grasslands, we found that grasses outperformed forbs in paired pots despite no differences when grown alone. Plants were well-watered and harvested before shading could become problematic, therefore differences in performance likely resulted from below-ground interactions. Given the large role played by microbes in below-ground interactions, the lack of a relationship between grass and forb performance in pairs, and the relationship between microbial abundance and plant performance in pairs, we speculate that invasive grass dominance over native forbs here is partially mediated by microbes. Invaded grassland soils tend to be depleted in plant available nitrogen due to the nutrient-demanding nature of invasive grasses [32,62,63], but with increased microbial activity and faster nitrogen cycling [24,31,63]. These microbe-driven changes may selectively benefit fast-growing grasses over natives, especially in nutrient-poor serpentine soils where added nitrogen has been found to preferentially aid invasives [51, 65]. Invasive grasses may further benefit from decreased microbial diversity, which has been linked to both decreased forb nitrogen uptake and increased grass nitrogen uptake [66]. Beyond competition for nutrients, grasses and forbs may also indirectly compete through recruitment of microbes harmful to natives [67] and/or helpful to invasives [36]. Other studies in serpentine grasslands have shown that soil primed with invasive grass negatively affected native forb growth due to changes in the microbial community [68, 69]. Although future work should be conducted to tease apart the role of plant density on microbial shifts and to understand the degree to which microbial shifts drive competitive outcomes in this system, our results add to a body of evidence that the invasion of grasses alters the microbiome and suggest that their dominance may be linked to these shifts.

Plant performance in pairs could not be explained by the performance of their neighbor. Instead, plant performance in pairs was tied to the abundance of key bacterial families. We found links between bacterial normalized abundance and plant biomass in grass-forb pairs for five of the six main differentially abundant families: three that were correlated with decreased forb performance: Burkholderiacae (marginal), Clostridacaeae_1 (marginal), and Fibrobacteraceae, three that were correlated with increased grass performance: Burkholderiaceae, Clostridiaceae_1 (marginal), and Veillonellaceae (marginal), and one that was correlated with both decreased grass performance and increased forb performance: Methylophilaceae.

Methylophilaceae have been found in soils with high heavy metal content and include microbes critical for heavy metal attenuation and immobilization [70, 71]. Members of Methylophilaceae also form beneficial symbioses with plants [72–74]. This family may include important taxa for serpentine-adapted species with roles in heavy metal attenuation and plant growth [73]. Forbs contributed a higher percentage of ASVs from this family to the joint grass-forb rhizosphere microbiome, further supporting close associations between native forbs and this group. If interactions with invasive grasses are driving declines in this family as our results suggest, grass dominance in this system may be partially due to a decrease in locally adapted microbes similar to Cavalieri et al. [30].

Grasses were likely driving higher normalized abundances of Fibrobacteraceae, Veillonellaceae, and Clostridiaceae_1, as they contributed a higher percentage of ASVs from these families to the joint rhizosphere microbiome in pairs. Although Fibrobacteraceae is typically lower in abundance in soils with high heavy metal content [71], this family is also positively associated with invasive grass dominance [75]. Further, all three families are known for their cellulose-degrading properties [76–78], and therefore may contain important taxa for organic matter degradation [79]. Both Fibrobacteraceae and Clostridiaceae_1 contain known nitrogen-fixers [80–82]. If taxa in these families are increasing nutrient availability, they could be disproportionately helping fast-growing invasive annual grasses [51, 65].

Burkholderiaceae are considered keystone members in grasslands as endophytes or pathogens [83–86] and are known for their antimicrobial properties [87–89], nitrogen-fixing abilities [80, 90], and competitive dominance, especially in N-limited environments [91]. Some members are even known to suppress fungal pathogens [92, 93]. It is possible that the Burkholderiacae observed in pairs here are either providing positive functional benefits to grasses, or are less harmful to grasses than forbs as pathogens.

In contrast to our bacterial findings, we found no significant differences in fungal communities across treatments, either in terms of structure or diversity. This suggests loose fungal associations with plant hosts, possibly due to functional redundancy of fungi across large geographic scales [94]. Within annual grasslands, however, there is evidence of invader-driven fungal shifts [61] and increases in fungal relative abundance [95]. Our limited fungal findings may indicate that fungi are generally less important than bacteria for competitive outcomes between plants [96]. Alternatively, fungi may be important, but complex fungal-fungal interactions at the community level may cancel out fitness effects on plants by beneficial and detrimental fungi [97]. In addition, experimental constraints such as unaccounted fungal regional source pools (such as airborne local spores or local watersheds [94]), plant age and development at time of sampling [16], and methodology (e.g. sampling rhizosphere here instead of root endophytes (e.g. Emam et al. [98]) or fungi in the rhizoplane (e.g. Edwards et al. [1])), may have prevented the observation of interactions between the fungal community and plant competition. *Conclusions*

Native annual forbs are host to a community of microbes distinct from, and more diverse than, those of their invasive competitors, but these close associations may be disrupted by invasive grasses. The joint bacterial rhizosphere of invasive grasses and native forbs differed from those of plants grown alone, with grasses contributing more to the abundance of ASVs from families that were linked to decreases in forb performance and/or increased in grass performance. Moreover, three of these families were higher in abundance in joint microbiomes relative to those of forbs alone. Forbs on the other hand contributed more to the abundance of ASVs from only one family that was associated with increased forb performance and decreased grass performance, but this family declined in abundance in the joint rhizosphere relative to forbs. Given the correlative nature of our study, more research is needed to understand the role of the microbiome in invasive grass dominance over native forbs and to clarify whether the observed changes in plant performance were indeed driven by these candidate bacterial families. Regardless, our study highlights the importance of considering the microbiome in ecosystems facing dominance by invasive species.

## Declarations

### Availability of data and material

The datasets and scripts supporting the conclusions of this article are available in the GitHub repository, DOI:10.5281/zenodo.4422181. The sequence reads generated for this 16S rRNA gene and ITS region amplicon project were deposited at Genbank under accession no. PRJNA666893.

### Code availability

No custom software was used. All R scripts used to process the data are available in the GitHub repository, DOI:10.5281/zenodo.4422181

### Authors’ contributions

MLL, CLE, and HK conceived of, planned, and carried out the experiment. MLL and CLE processed and analyzed the data and wrote the paper. All authors contributed to the final version of the manuscript.

### Ethics approval

Not applicable.

### Consent to participate

Not applicable.

### Consent for publication

Not applicable.

## Acknowledgements

We would like to thank undergraduate intern, Vyvy Ha, with their help on this project, including processing plant biomass and assisting with shade-house related activities. Finally, we would like to thank Susan P. Harrison, Andrew M. Latimer, Jonathan A. Eisen, Lauren M. Hallett, and Jennifer R. Gremer for useful conversations and suggestions related to this work, providing lab resources and space and contributing funds to this project.

