## Supplemental Methods, Figures & Tables for "Invasive grass dominance over native forbs is linked to shifts in the bacterial rhizosphere microbiome"

*Experimental set-up*

In spring and summer 2017, we collected soil and seed from an annual grassland at the University of California McLaughlin Natural Reserve [1] in the Inner North Coast Range (N 38°52', W 122°26'). The site has a Mediterranean climate with cool, wet winters (mean annual precipitation 2000-2019 of 738 mm [2]) and dry, hot summers. Soil was collected from an area of the reserve with deep, finely-textured serpentine soils and seeds were collected nearby from six common native annual forb species (Fig. 1a; *Lasthenia californica, Clarkia purpurea, Agoseris heterophylla, Calycadenia pauciflora, Hemizonia congesta,* and *Plantago erecta)* and three dominant invasive annual grasses (Fig. 1b; *Avena fatua, Taeniatherum caput-medusae,* and *Bromus hordeaceus*). In December 2017, seeds were sown into physan-washed pots with a mixture of 65% field-collected soil and 35% sterile autoclaved sand to improve drainage. Each species was grown alone (one individual per pot, five replicate pots per species) to assess the microbial communities of individual plant species. To assess the effects of competition between natives and invasives, each forb species was grown with each grass species in a pairwise factorial design for a total of 18 different species combinations (one individual of each grass and forb per pot, five replicate pots per combination). Pots were placed in a shade-house open to natural temperature variation in the UC Davis Orchard Park Greenhouse and soil moisture was maintained with an automatic drip irrigation system (Fig. 1c).

*Sample collection*

In April 2018, we sampled rhizosphere microbial communities by gently shaking off soil from plant roots, then submerging the roots in autoclaved nanopure water in a 50 mL conical tube and vortexing to obtain the rhizosphere soil similar to Edwards et al. [3]. We sampled rhizospheres prior to flowering to minimize differences in composition due to developmental stage [4, 5]. Conical tubes were then centrifuged at 4000 g for 1 minute to obtain rhizosphere soil pellets. In grass-forb pairs, we sampled the joint rhizosphere microbiome, similar to Sun et al. [6], as roots were completely interwoven.

*Molecular methods and sequence generation*

We randomized samples using a random number generator prior to DNA extraction. We then extracted the DNA from samples using the MoBio PowerSoil DNA Extraction kit (MO BIO Laboratories, Inc., Carlsbad, CA, USA) with one minor modification to manufacturer's instructions: samples were bead beat on the “homogenize” setting for 1 minute during step 5. Samples were then sent to the Integrated Microbiome Resource (IMR) at Dalhousie University to amplify and sequence the 16S rRNA gene using the 515FB-806RB primer set [7, 8] and ITS2 region using the ITS86F-ITS4R primer set [9]. The amplification and sequencing process is described in Comeau et al. [10]. In addition to 129 rhizosphere experimental samples, we sent five subsamples of the background field-collected soil-sand mix, three replicates each of a positive technological control using DNA extracted from the ZymoBIOMICS Microbial Community Standard (Zymo Research Inc., Irvine, CA, USA), and a negative technological control where no sample was added during DNA extraction to the IMR for sequencing. The sequence reads generated for this 16S rRNA gene and ITS region amplicon project were deposited at Genbank under accession no. PRJNA666893.

*Sequence processing*

Sequencing primers were removed using the Cutadapt (v. 2.3) program and the resulting sequences were then processed using the DADA2 workflow in R [11–13]. 16S rRNA gene amplicon sequences were trimmed at 280 base pairs for forward reads and 250 base pairs for reverse reads based on their error profiles to maintain a quality score above 10. ITS region amplicon sequences were not trimmed to preserve the ability to merge reads due to the inherent length variation of the ITS region. Both 16S rRNA gene and ITS region amplicon reads were then truncated at the first quality score of 2 and reads with an expected error greater than 2 were removed. We then merged paired end reads, created an Amplicon Sequence Variant table (ASV), and identified chimeric reads using removeBimeraDenovo, which were subsequently removed prior to downstream analyses (~4.1% of sequence reads for 16S rRNA gene, ~0.93% for ITS region).

For 16S rRNA gene ASVs, we assigned taxonomy using the RDP Naive Bayesian Classifier algorithm and the SILVA high quality ribosomal RNA database v. 132 [14]. For ITS ASVs, taxonomy was inferred with the UNITE (v. 8.0) database modified to include a representative ITS amplicon sequence for each host plant, i.e., *H. congesta* (MF964056.1), *C. purpurea* (MG235764.1), *P. erecta* (AY101909.1), *A. fatua* (FJ794719.1), *L. californica* (AY043514.1), *A. heterophylla* (AY218965.1), *T. caput-medusae* (AJ608153.1), *B. hordeaceus* (KP987344.1) [15, 16]. ITS-x (v. 1.1.1) was then run on the unique ITS ASVs to pull out only sequences that had a fungal ITS region based on kingdom-specific hidden Markov models [17]. ITS-x was unable to detect a fungal ITS region in 744 of the ASVs (~13.46% of ASVs). These ASVs were thus considered putatively non-fungal and were removed from downstream analysis.

Decontam's prevalence method was used to identify putative contaminants with a threshold of 0.5 which will identify sequences that have a higher prevalence in negative controls than in true samples [18]. This method identified 55 contaminants in the 16S rRNA gene data and 32 in the ITS amplicon data which were then removed from the analysis.

For the 16S rRNA gene dataset, unique ASVs were aligned using the MAFFT Alignment Program (v 7.402) on XSEDE with standard parameters through the CIPRES Science Gateway V. 3.3 (https://www.phylo.org/) [19, 20]. We then built a phylogenetic tree using FastreeMP on XSEDE with standard parameters also through CIPRES [21] and rooted the tree using an archaeal outgroup. We then created a phyloseq object from the resulting 16S ASV table, Silva taxonomy table, phylogeny, and mapping file using the phyloseq package in R [22]. Sequences that were taxonomically assigned to chloroplasts and mitochondria were then removed. Samples were rarefied to 9434 reads per sample for questions 1-3.

For the ITS region data, we created a phyloseq object from the ITS ASV table, UNITE taxonomy table, and mapping file using the phyloseq package in R. Samples were rarefied to 7557 reads per sample for questions 1-3.

The datasets and scripts supporting the conclusions of this article are available in the GitHub repository, DOI:10.5281/zenodo.4422181

**Supplemental Tables and Figures:**

**Table S1.** Untransformed mean relative abundances of the 19 differentially abundant bacterial families and 3 differentially abundant fungal families

**Table S2.** Regression results for correlations of plant log response ratios and family normalized abundance

**Figure S1.** Fungal rhizosphere community structures do not vary between grasses and forbs, and do not shift during competition

**Figure S2.** Fungal Shannon diversity does not differ between grasses and forbs and do not shift during competition

**Figure S3.** Rhizospheres differ structurally from background soil in both (a) bacterial and (b) fungal communities

**Figure S4.** Bacterial rhizosphere community structure did not vary between forb species, but did vary between grass species

**Figure S5.** Fungal rhizosphere community structure did not vary between forb or grass species

**Figure S6.** Differentially abundant (a) bacterial and (b) fungal families across treatments

**Figure S7.** Normalized abundance of remaining 13 bacterial families across treatments

**Figure S8.** Normalized abundance of fungal families across treatments

**Figure S9.** Bacterial family relationships to plant performance for remaining 13 families

**Figure S10.** Predicted sources for ASVs in differentially abundant bacterial families

**Figure S11.** Fungal family relationships to plant biomass in pairs

**Figure S12.** Majority of ASVs in differentially abundant fungal families correlated with competition were sourced from grasses

**Table S1.** Untransformed mean relative abundance and standard error of the 19 differentially abundant bacterial families and 3 differentially abundant fungal families.

| **Rhizosphere** | **Family** | **Mean Relative Abundance (%)** | **Standard Error** |
|---|---|---|---|
| Bacterial | Acetobacteraceae | 0.72 | 0.03 |
| | Azospirillaceae | 3.24 | 0.12 |
| | Beijerinckiaceae | 2.63 | 0.07 |
| | Burkholderiaceae | 11.69 | 0.39 |
| | Clostridiaceae_1 | 0.13 | 0.02 |
| | Cyclobacteriaceae | 0.11 | 0.01 |
| | Fibrobacteraceae | 0.32 | 0.02 |
| | Flavobacteriaceae | 1.78 | 0.14 |
| | Geodermatophilaceae | 1.93 | 0.10 |
| | Methylophilaceae | 0.35 | 0.03 |
| | Microbacteriaceae | 0.17 | 0.01 |
| | Nitrospiraceae | 0.19 | 0.01 |
| | Propionibacteriaceae | 0.79 | 0.04 |
| | Rhodocyclaceae | 0.75 | 0.16 |
| | Rubritaleaceae | 0.08 | 0.01 |
| | Steroidobacteraceae | 0.65 | 0.04 |
| | Streptomycetaceae | 0.18 | 0.02 |
| | Veillonellaceae | 0.12 | 0.03 |
| | Weeksellaceae | 0.08 | 0.01 |
| Fungal | Ceratobasidiaceae | 2.64 | 0.60 |
| | Tubeufiaceae | 0.23 | 0.09 |
| | Unclassified Sebacinales | 0.60 | 0.16 |

**Table S2.** Regression results comparing normalized abundance of each differentially abundant family to biomass of grasses and forbs in paired pots. Families in bold indicate those that were found to vary in abundance between grasses and forbs, and between grass-forb pairs and either grasses or forbs. Bolded *p*-values indicate significance ($p < 0.050$), *p*-values in parentheses indicate marginal significance ($p < 0.10$). See Figs. 6, S10, S12.

| Rhizosphere | Family | Group | Estimate | SE | *p*-value |
|---|---|---|---|---|---|
| Bacterial | Acetobacteraceae | forb | 0.01 | 0.21 | 0.972 |
| | | grass | 0.04 | 0.20 | 0.825 |
| | Azospirillaceae | forb | -0.02 | 0.16 | 0.909 |
| | | grass | 0.09 | 0.15 | 0.575 |
| | Beijerinckiaceae | forb | 0.10 | 0.24 | 0.682 |
| | | grass | 0.15 | 0.23 | 0.524 |
| | **Burkholderiaceae** | forb | -0.26 | 0.15 | (0.091) |
| | | grass | 0.36 | 0.16 | **0.026** |
| | **Clostridiaceae_1** | forb | -0.29 | 0.17 | (0.097) |
| | | grass | 0.31 | 0.17 | (0.066) |
| | Cyclobacteriaceae | forb | 0.13 | 0.28 | 0.635 |
| | | grass | -0.42 | 0.24 | (0.083) |
| | **Fibrobacteraceae** | forb | -0.40 | 0.19 | **0.042** |
| | | grass | -0.10 | 0.18 | 0.601 |
| | Flavobacteriaceae | forb | -0.10 | 0.09 | 0.271 |
| | | grass | 0.09 | 0.08 | 0.287 |
| | Geodermatophilaceae | forb | 0.07 | 0.14 | 0.610 |
| | | grass | 0.02 | 0.13 | 0.856 |
| | **Methylophilaceae** | forb | 0.40 | 0.14 | **0.007** |
| | | grass | -0.31 | 0.14 | **0.029** |
| | Microbacteriaceae | forb | 0.18 | 0.24 | 0.458 |
| | | grass | 0.00 | 0.23 | 0.989 |
| | Nitrospiraceae | forb | -0.05 | 0.34 | 0.879 |
| | | grass | 0.06 | 0.31 | 0.833 |
| | Propionibacteriaceae | forb | 0.04 | 0.16 | 0.781 |
| | | grass | 0.14 | 0.15 | 0.350 |
| | Rhodocyclaceae | forb | -0.08 | 0.06 | 0.211 |
| | | grass | 0.03 | 0.06 | 0.594 |
| | Rubritaleaceae | forb | -0.37 | 0.26 | 0.153 |

| | | | | | |
|---|---|---|---|---|---|
| | | grass | 0.45 | 0.24 | (0.067) |
| | Steroidobacteraceae | forb | -0.12 | 0.26 | 0.642 |
| | | grass | -0.40 | 0.25 | 0.109 |
| | Streptomycetaceae | forb | -0.03 | 0.17 | 0.883 |
| | | grass | 0.06 | 0.17 | 0.728 |
| | **Veillonellaceae** | forb | -0.11 | 0.15 | 0.453 |
| | | grass | 0.24 | 0.14 | (0.090) |
| | Weeksellaceae | forb | -0.11 | 0.17 | 0.516 |
| | | grass | 0.35 | 0.16 | **0.029** |
| Fungal | Ceratobasidiaceae | forb | -0.13 | 0.05 | **0.013** |
| | | grass | 0.22 | 0.05 | **<0.001** |
| | **Tubeufiaceae** | forb | 0.06 | 0.13 | 0.670 |
| | | grass | 0.01 | 0.13 | 0.959 |
| | Unclassified Sebacinales | forb | 0.01 | 0.03 | 0.777 |
| | | grass | 0.02 | 0.03 | 0.360 |

**Figure S1.** Fungal rhizosphere community structures do not vary between grasses and forbs, and do not shift during competition. Principal coordinates analysis (PCoA) visualization of Bray Curtis dissimilarities of fungal communities associated with the rhizosphere. Points in the ordination are colored and represented by shapes based on the rhizosphere of forbs grown alone (purple circles), grasses grown alone (green triangles) and grass-forb competition pairs (yellow squares).

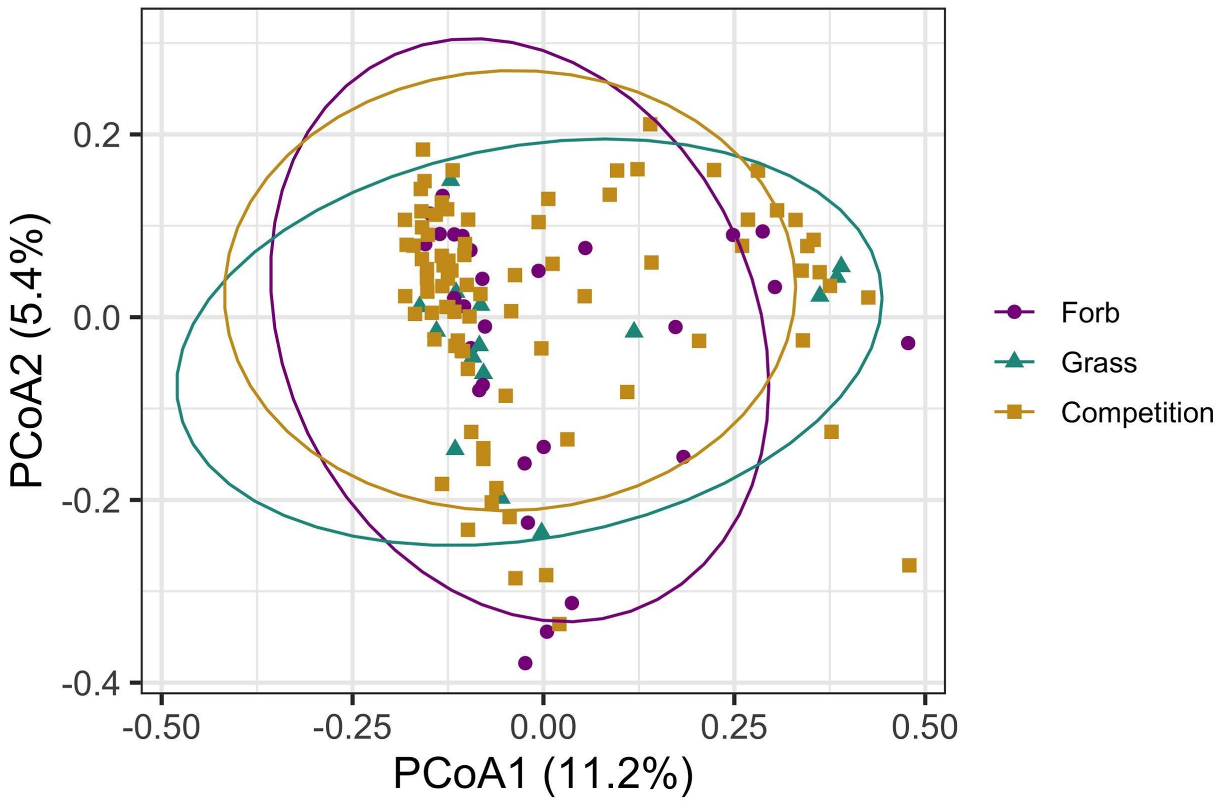

**Figure S2.** Fungal Shannon diversity does not differ between grasses and forbs and do not shift during competition. Shannon diversity was used to assess alpha diversity for fungal communities for each treatment (forbs grown alone, grasses grown alone, grass-forb competition pairs). Comparisons that are significantly different from each other are notated by different letters (e.g. a vs. b).

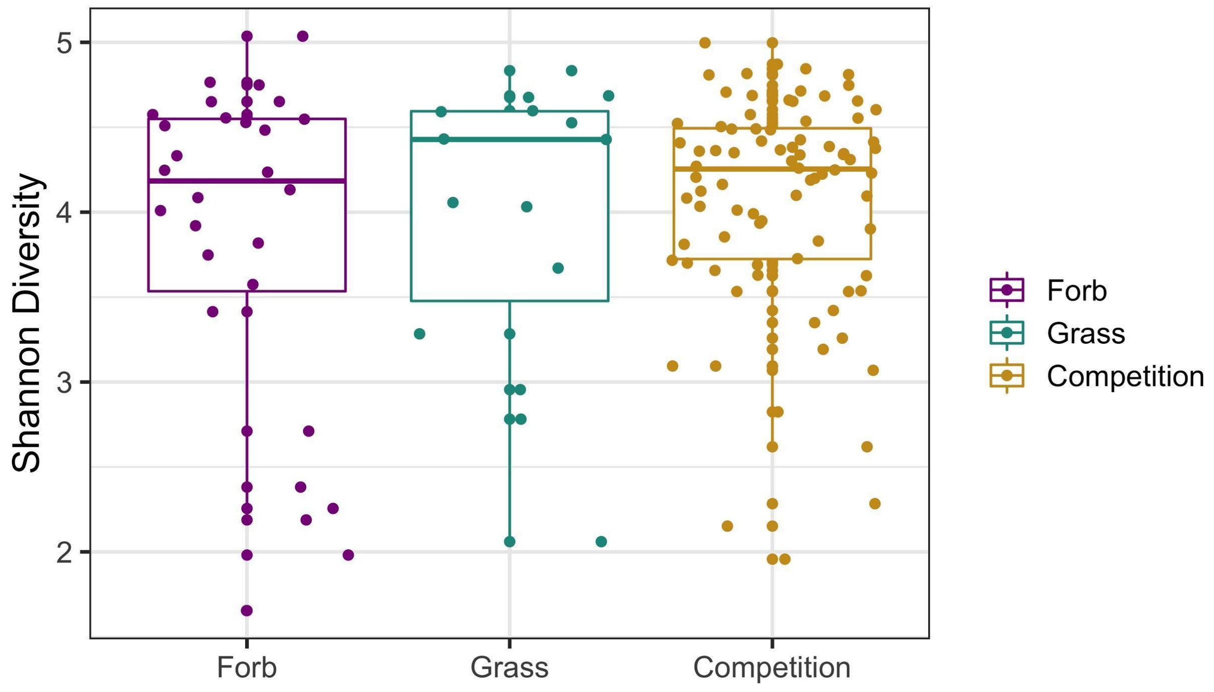

**Figure S3.** Rhizospheres differ structurally from background soil in both (a) bacterial and (b) fungal communities.

(a)

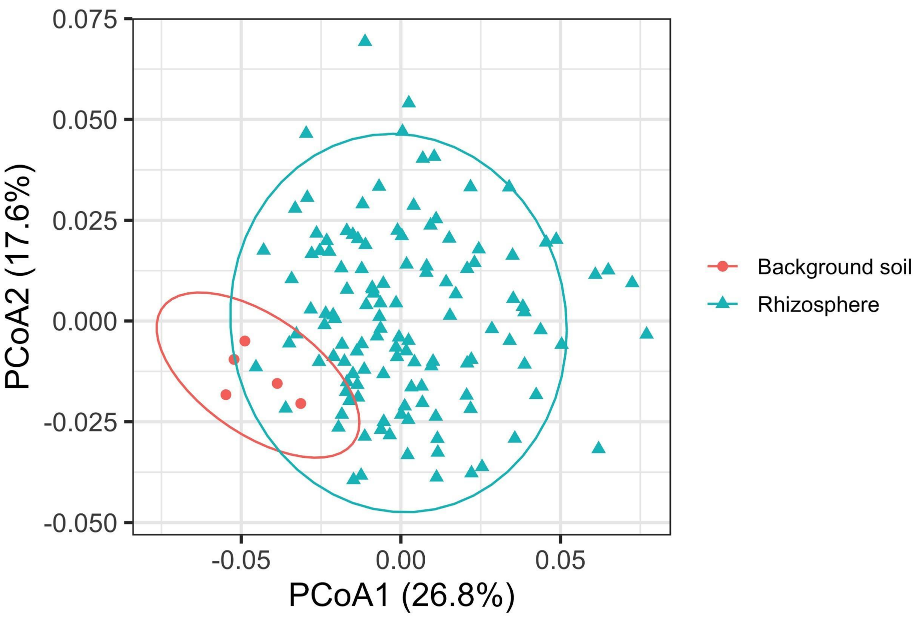


(b)

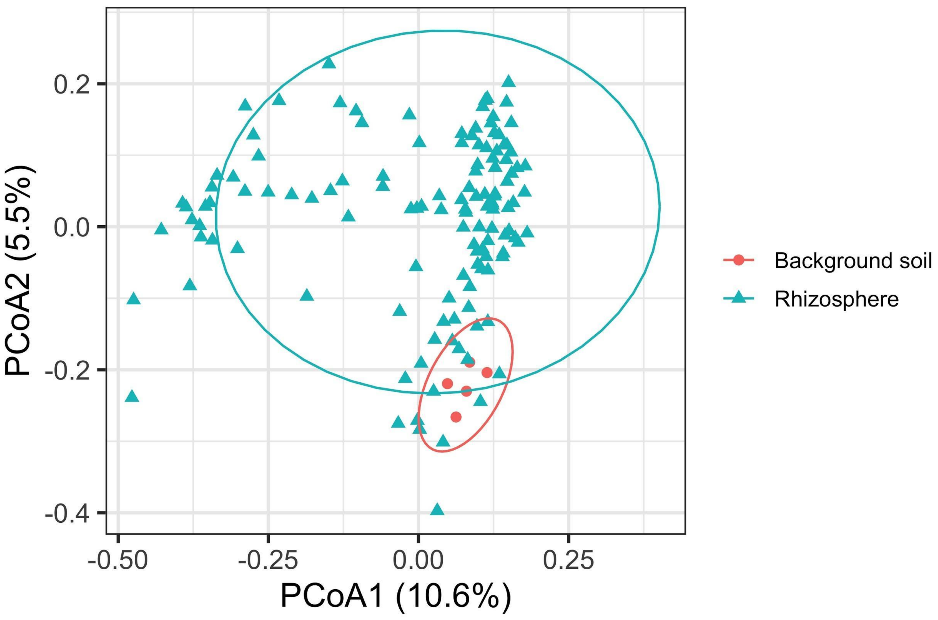

**Fig S4.** (a) Bacterial rhizosphere microbiomes did not vary between native forb species ($p$ = 0.069), but (b) did vary between invasive grass species ($p$ = 0.013), with the microbiomes of *T. caput-medusae* displaying different structure and composition from the other two grasses (*B. hordeaceus*: $p$ = 0.049; *A. fatua:* $p$ = 0.047). Principal coordinates analysis (PCoA) visualization of weighted UniFrac distances of bacterial communities associated with the rhizosphere. Points in the ordination are colored and represented by shapes based on the rhizosphere of different forb and grass species grown alone.

(a) Native forbs

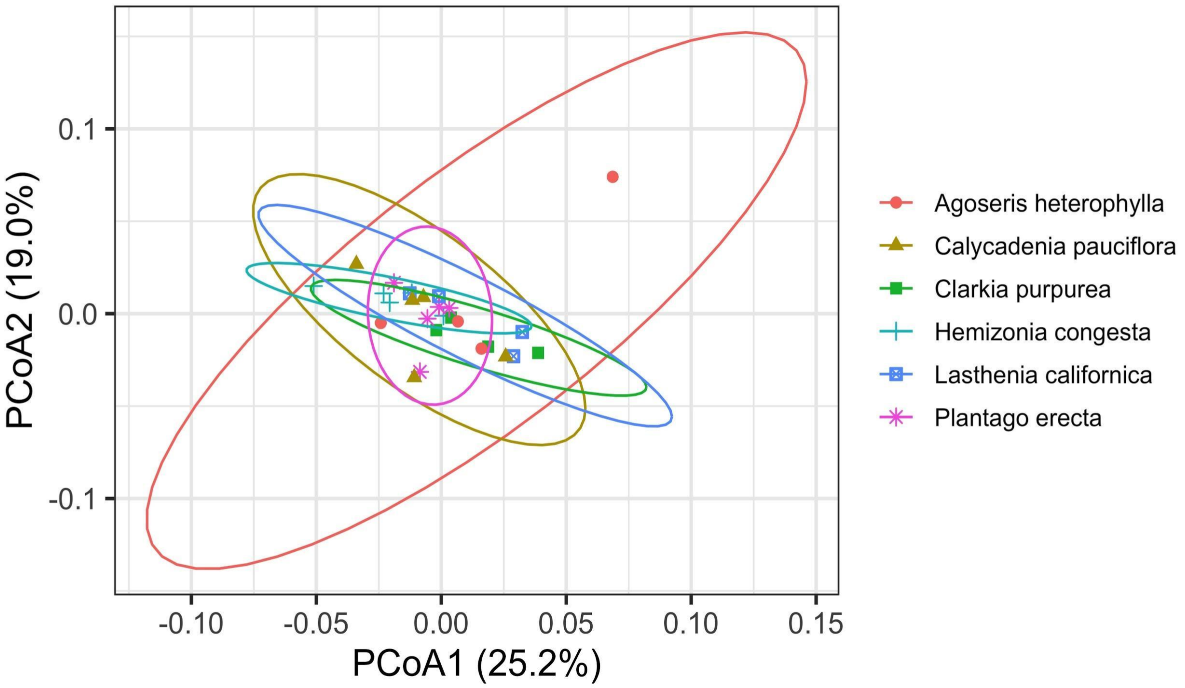

(b) Invasive grasses

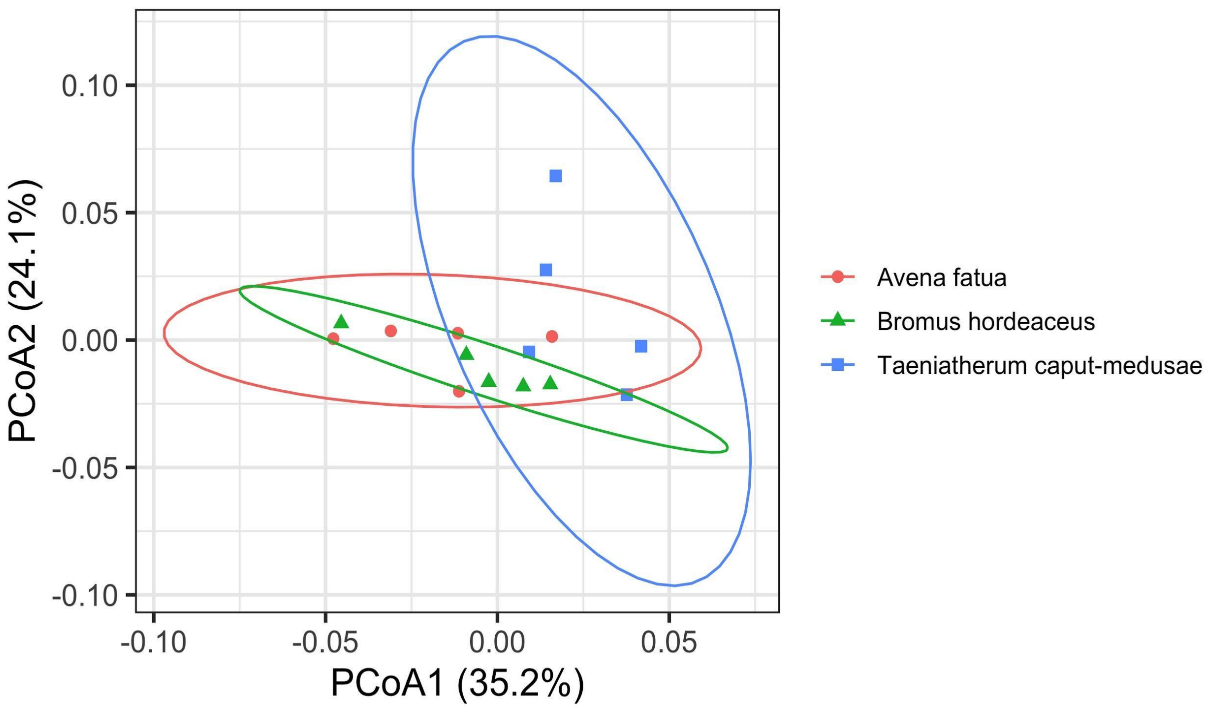

**Fig S5.** (a) Fungal rhizosphere microbiomes did not vary between native forb species ($p$ = 0.755) or (b) between invasive grass species ($p$ = 0.589). Principal coordinates analysis (PCoA) visualization of Bray Curtis dissimilarities of fungal communities associated with the rhizosphere. Points in the ordination are colored and represented by shapes based on the rhizosphere of different forb and grass species grown alone.

(a) Native forbs

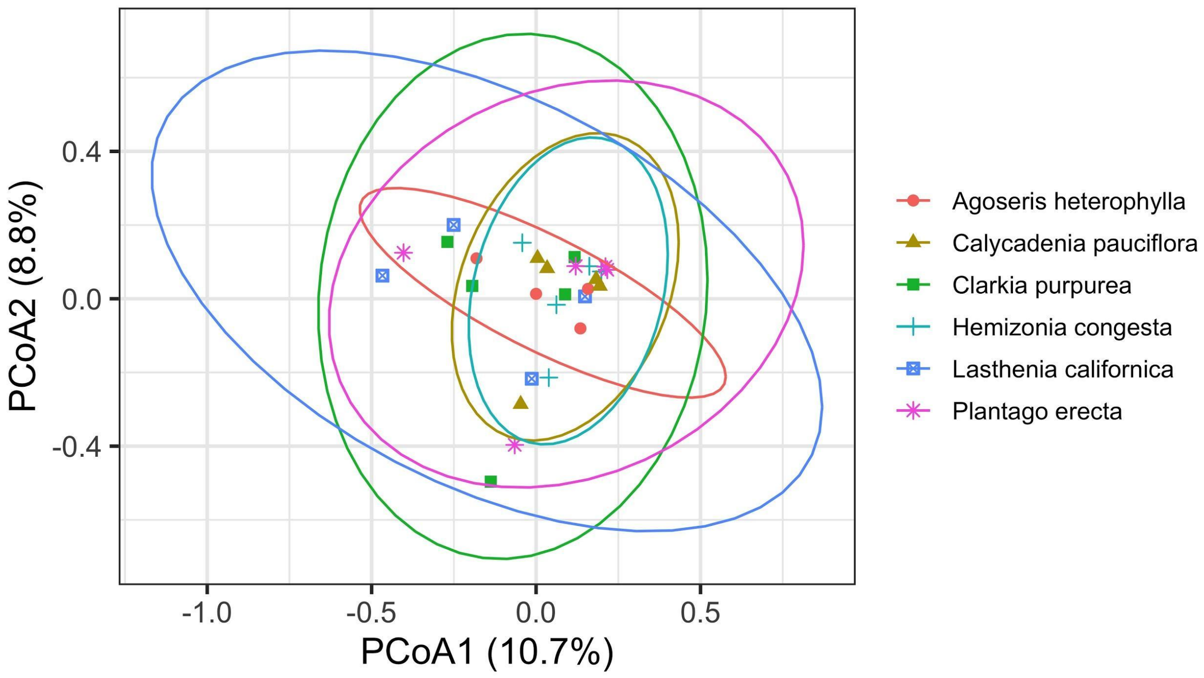

(b) Invasive grasses

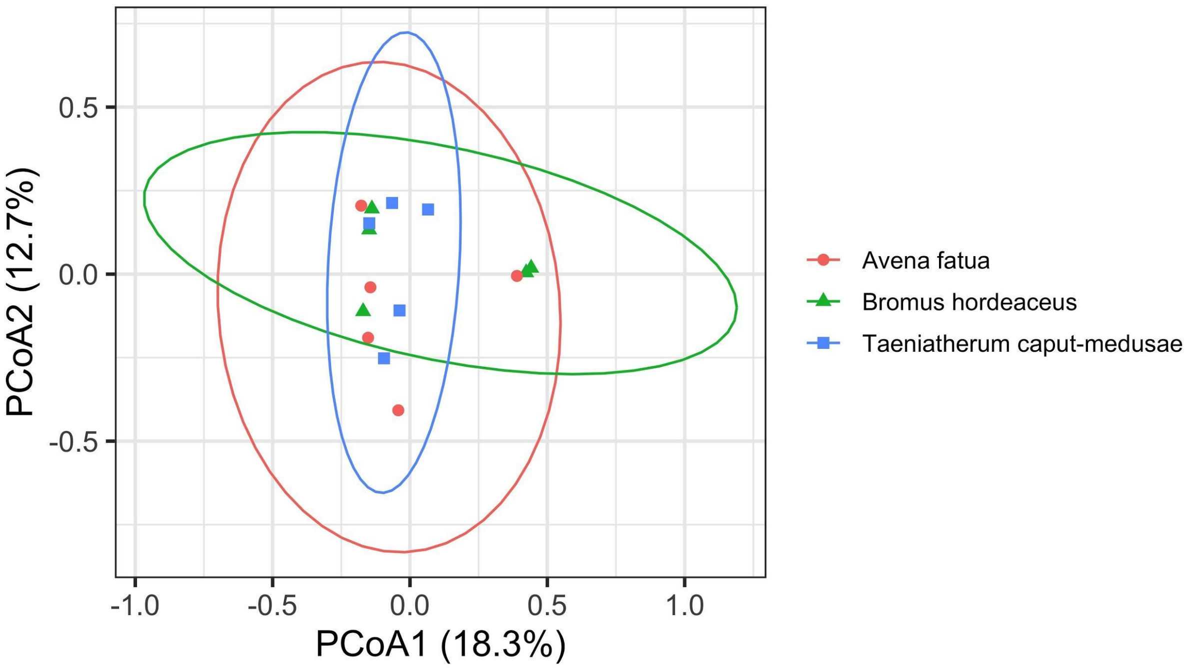

**Figure S6.** Differentially abundant families across treatments. Bacterial families (a) and fungal families (b) were identified whose abundance differed significantly between treatments (forbs grown alone, grasses grown alone, competition) using DESseq2. Each plot shows the log2 fold change of families which were differentially abundant between grass grown alone and forbs grown alone (left), grass-forb pairs (hereafter referred to as competition) and forbs grown alone (middle), and competition and grasses grown alone (right). A positive $\log_2$ fold change means the family was more abundant in the first treatment and a negative $\log_2$ fold change means the family was more abundant in the latter treatment.

(a) Bacterial

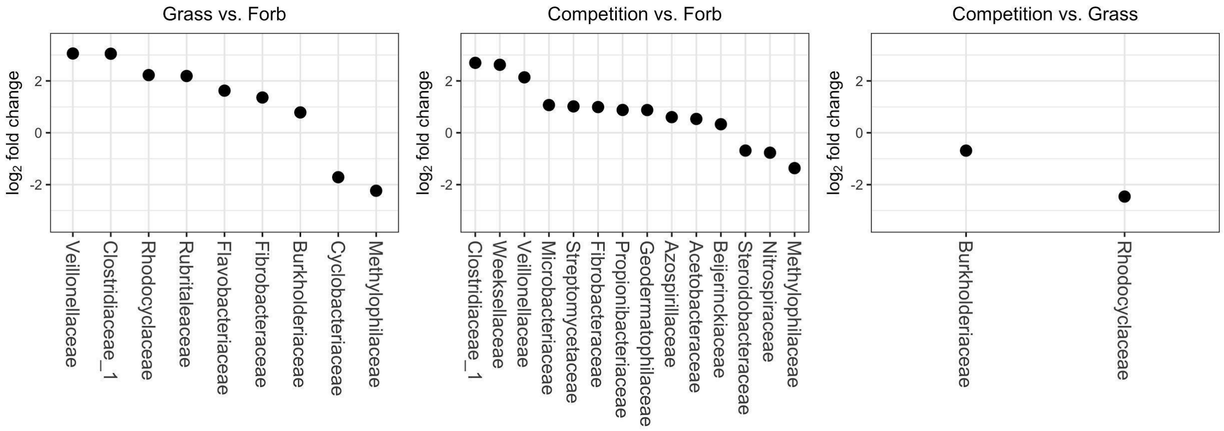


(b) Fungal

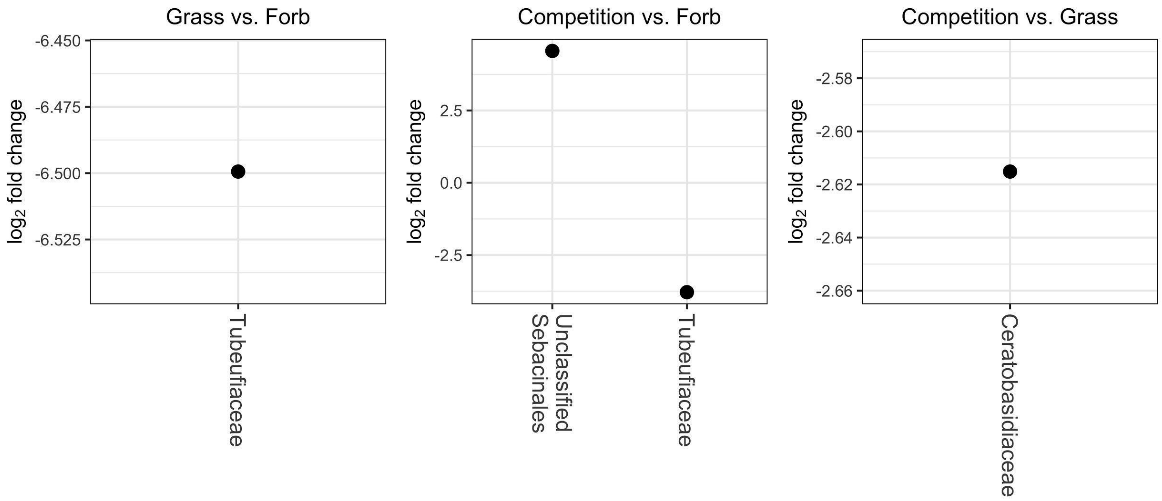

**Figure S7.** Normalized abundance of remaining 13 differentially abundant bacterial families. Using DESeq2, bacterial families were identified whose abundance differed significantly between treatments (forbs grown alone, grasses grown alone, forb-grass pairs). Plots here represent families that only varied between one combination of treatments. Each plot shows the normalized abundance of families for each treatment and comparisons that are significant from each other are notated by different letters (e.g., a vs. b).

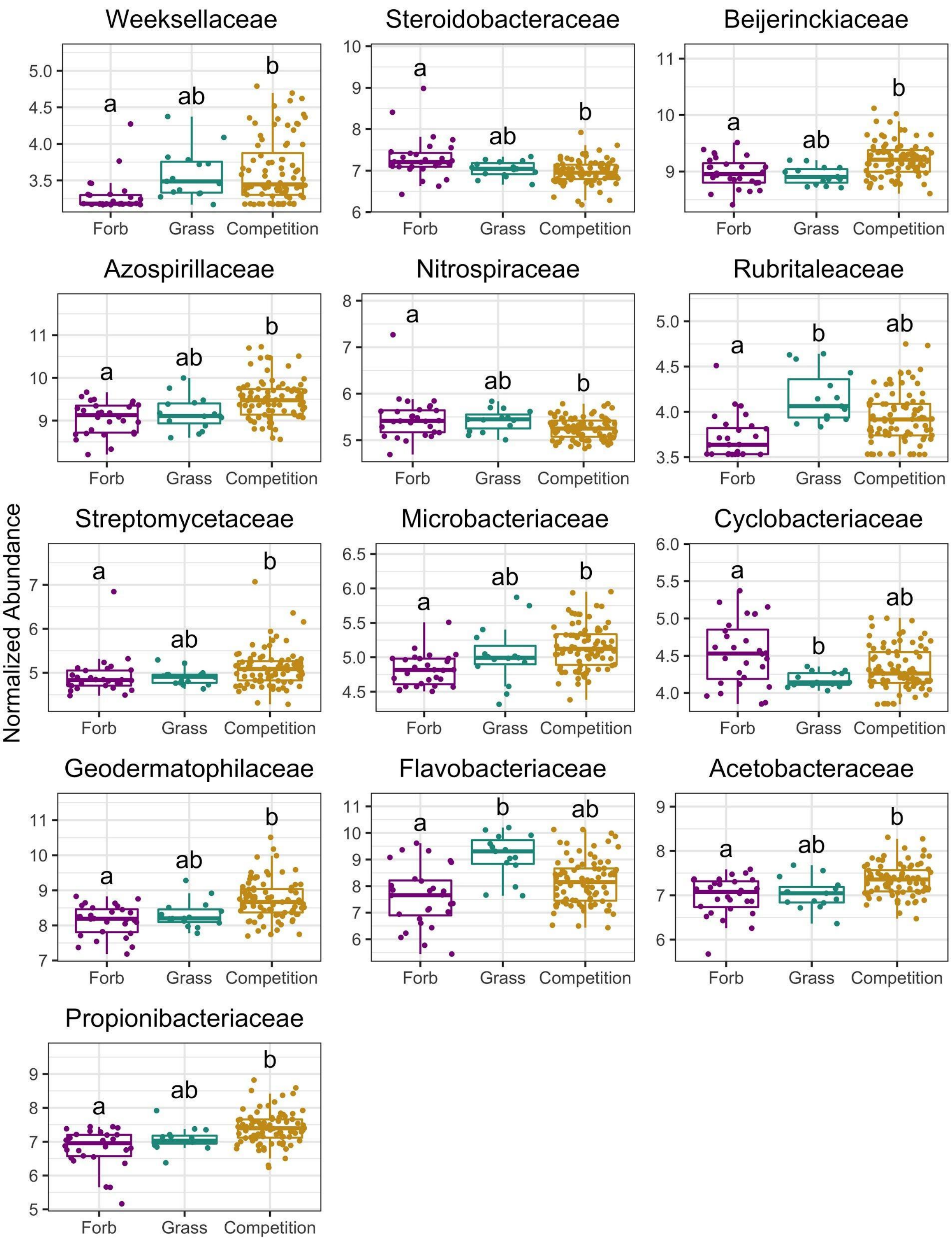
Normalized Abundance
Weeksellaceae
Steroidobacteraceae
Beijerinckiaceae
Azospirillaceae
Nitrospiraceae
Rubritaleaceae
Streptomycetaceae
Microbacteriaceae
Cyclobacteriaceae
Geodermatophilaceae
Flavobacteriaceae
Acetobacteraceae
Propionibacteriaceae
Forb
Grass
Competition
a
ab
b

**Figure S8.** Normalized abundance of fungal families across treatments. Using DESeq2, fungal families were identified whose abundance differed significantly between treatments (forbs grown alone, grasses grown alone, grown together in competition). Each plot shows the normalized abundance of families for each treatment and comparisons that are significant from each other are notated by different letters (e.g., a vs. b).

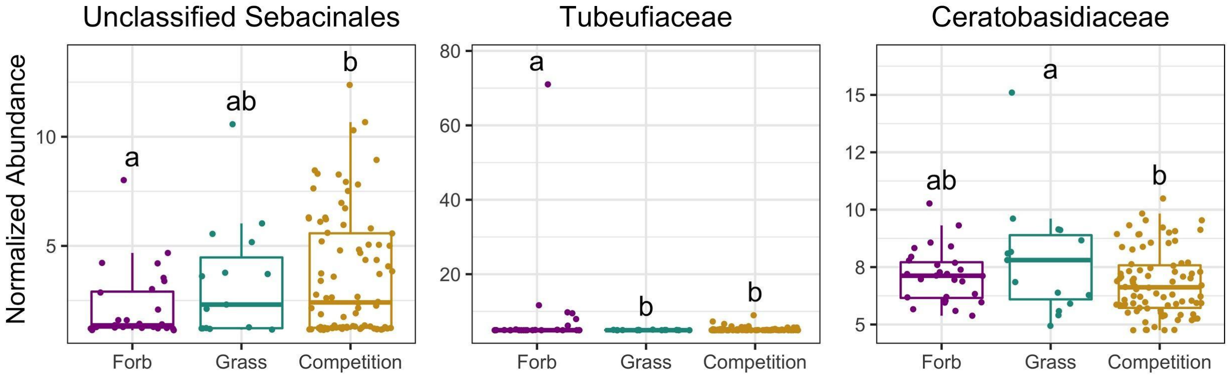

**Figure S9.** Normalized abundance of 13 bacterial families in relation to plant biomass in pairs for forbs (purple) or grasses (green). Plots here represent families that only varied between one combination of treatments. Solid lines indicate significance ($p < 0.05$), dashed lines indicate marginal significance ($p < 0.10$).

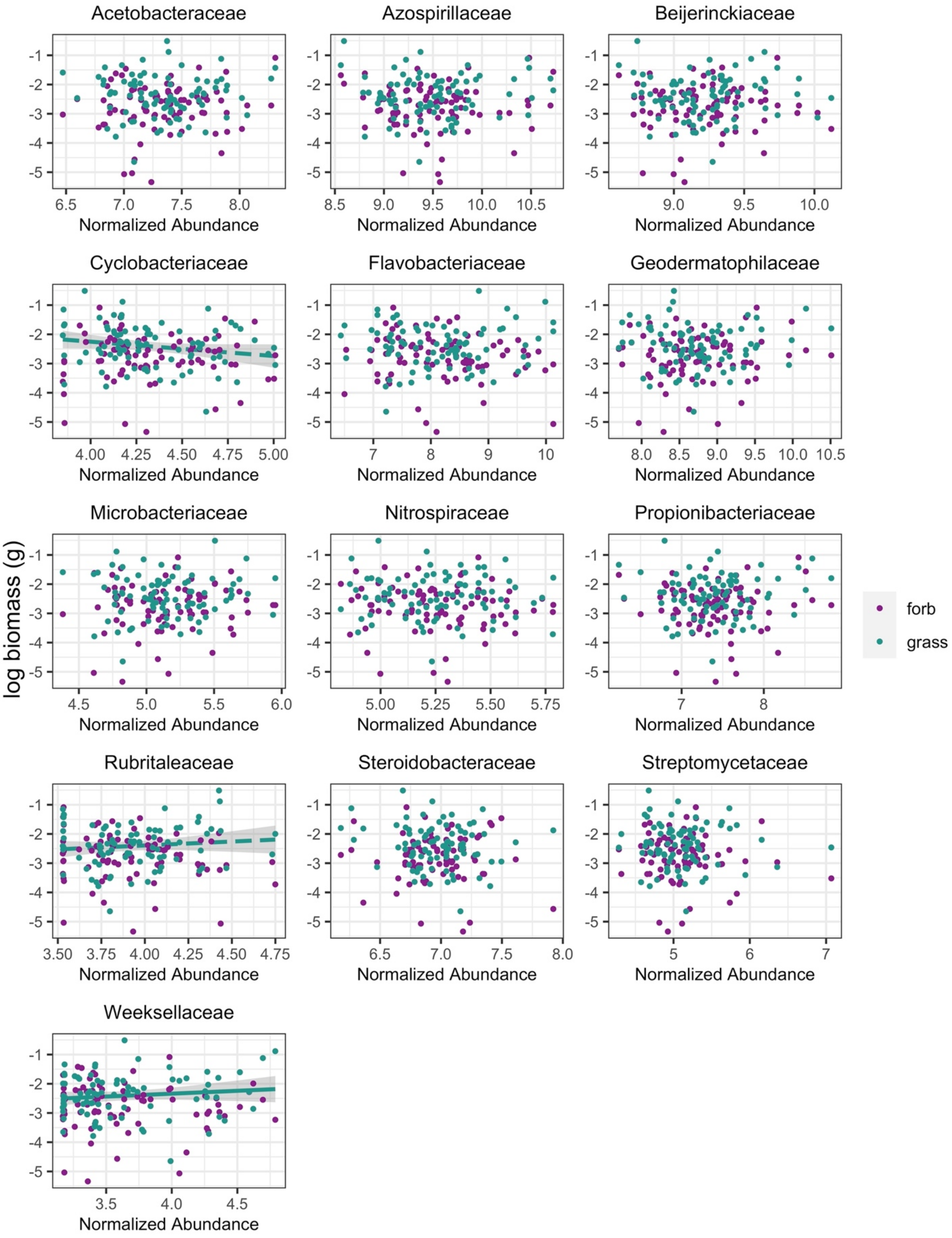

Acetobacteraceae
Azospirillaceae
Beijerinckiaceae
Cyclobacteriaceae
Flavobacteriaceae
Geodermatophilaceae
Microbacteriaceae
Nitrospiraceae
Propionibacteriaceae
Rubritaleaceae
Steroidobacteraceae
Streptomycetaceae
Weeksellaceae
log biomass (g)
Normalized Abundance
forb
grass

**Figure S10.** Predicted sources for ASVs in bacterial families that did not vary between both grasses and forbs and between pairs and individuals. SourceTracker was used to predict whether ASVs in competition treatments originated from grasses (dark green), forbs (purple), the background soil mix used in the experiment (blue), or unknown habitats (light green) and then the results were summarized by grouping ASVs by taxonomic family.

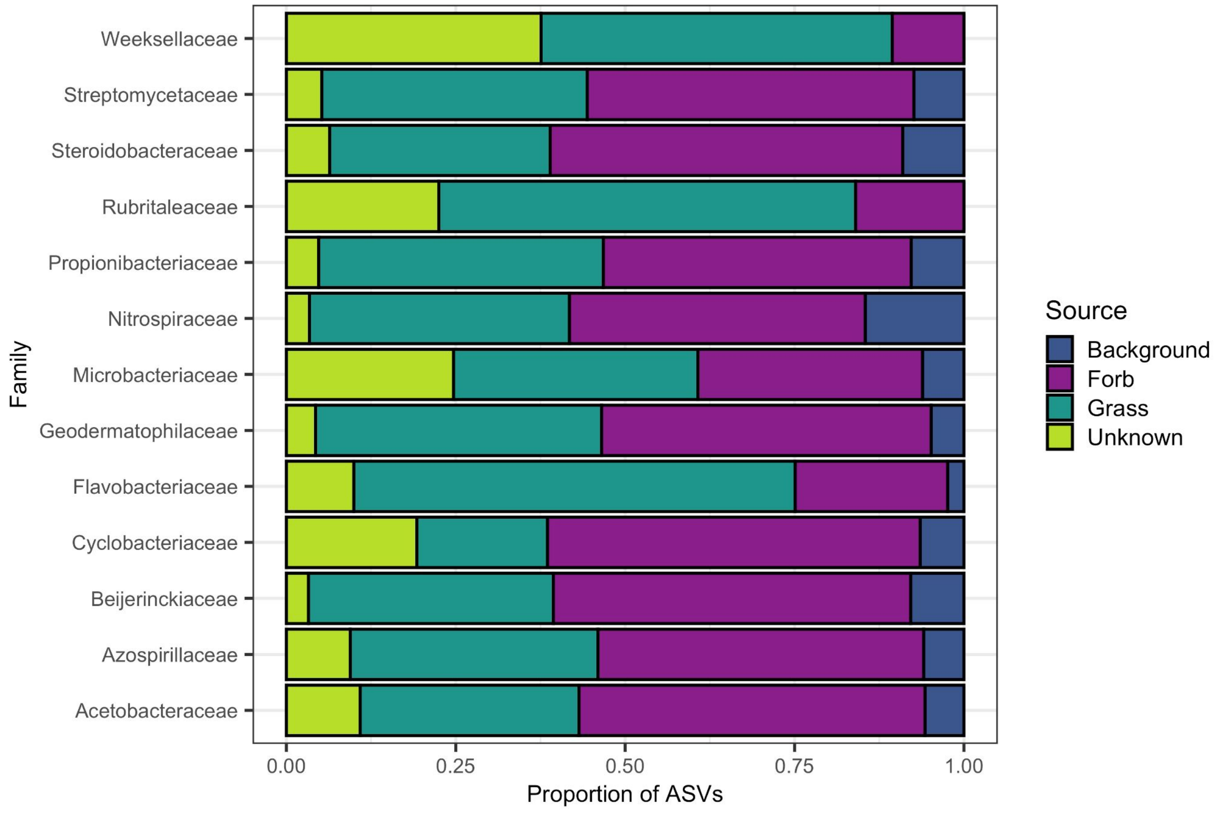

**Figure S11.** Fungal family relationships to plant biomass in pairs. Linear mixed effects models assessing the correlation of normalized abundance of fungal families to plant biomass for forbs (purple) or grasses (green). Solid line indicates significance ($p < 0.05$).

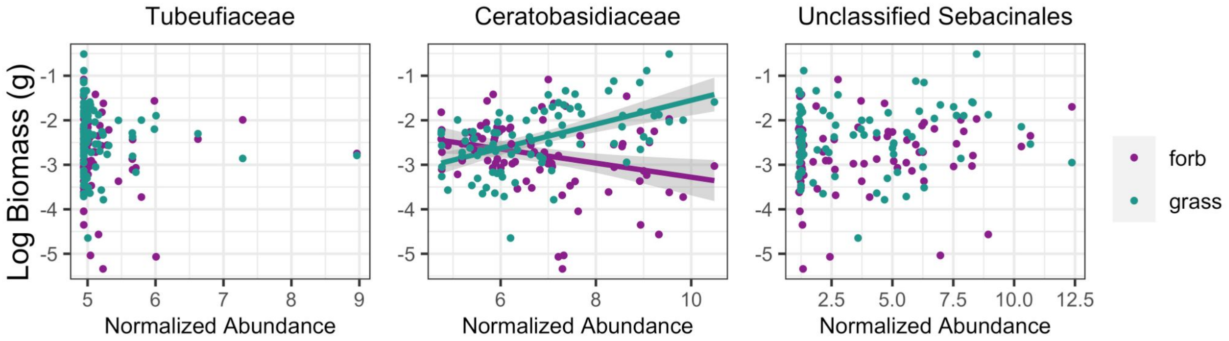

**Figure S12.** Proportion of ASVs predicted to colonize from each source for each of the three differentially abundant fungal families. SourceTracker was used to predict whether ASVs in competition treatments originated from grasses (dark green), forbs (purple), the background soil mix used in the experiment (blue), or unknown habitats (light green) and then the results were summarized by grouping ASVs by taxonomic family. The majority of ASVs in Ceratobasidiaceae, the only differentially abundant fungal family that was correlated with competition, were sourced from grass.

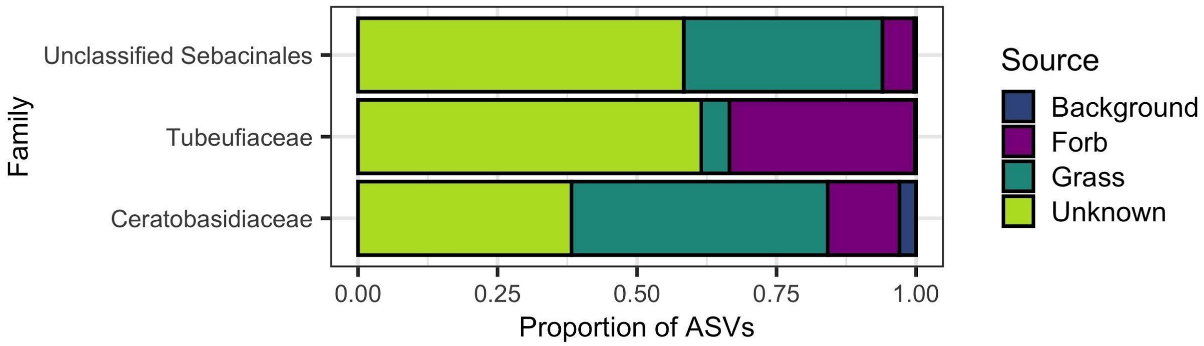
